# Minimizing the *ex vivo* confounds of cell-isolation techniques on transcriptomic -profiles of purified microglia

**DOI:** 10.1101/2021.07.15.452509

**Authors:** Sarah R. Ocañas, Kevin D. Pham, Harris E. Blankenship, Adeline H. Machalinski, Ana J. Chucair-Elliott, Willard M. Freeman

## Abstract

Modern molecular neuroscience studies require analysis of specific cellular populations derived from brain tissue samples to disambiguate cell-type specific events. This is particularly true in the analysis of minority glial populations in the brain, such as microglia, which may be obscured in whole tissue analyses. Microglia have central functions in development, aging, and neurodegeneration and are a current focus of neuroscience research. A long-standing concern for glial biologists using *in vivo* models is whether cell isolation from CNS tissue could introduce *ex vivo* artifacts in microglia, which respond quickly to changes in the environment. Mouse microglia were purified by magnetic-activated cell sorting (MACS), as well as cytometer- and cartridge-based fluorescence-activated cell sorting (FACS) approaches to compare and contrast performance. The Cx3cr1-NuTRAP mouse model was used here to provide an endogenous fluorescent microglial marker and a microglial-specific translatome profile as a baseline comparison lacking cell isolation artifacts. All methods performed similarly for microglial purity with main differences being in cell yield and time of isolation. *Ex vivo* activation signatures occurred principally during the initial tissue dissociation and cell preparation and not the microglial cell sorting. Utilizing transcriptional and translational inhibitors during the cell preparation prevented the activational phenotype. These data demonstrate that a variety of microglial isolation approaches can be used, depending on experimental needs, and that inhibitor cocktails are effective at reducing cell preparation artifacts.

**Table of Contents Image:** 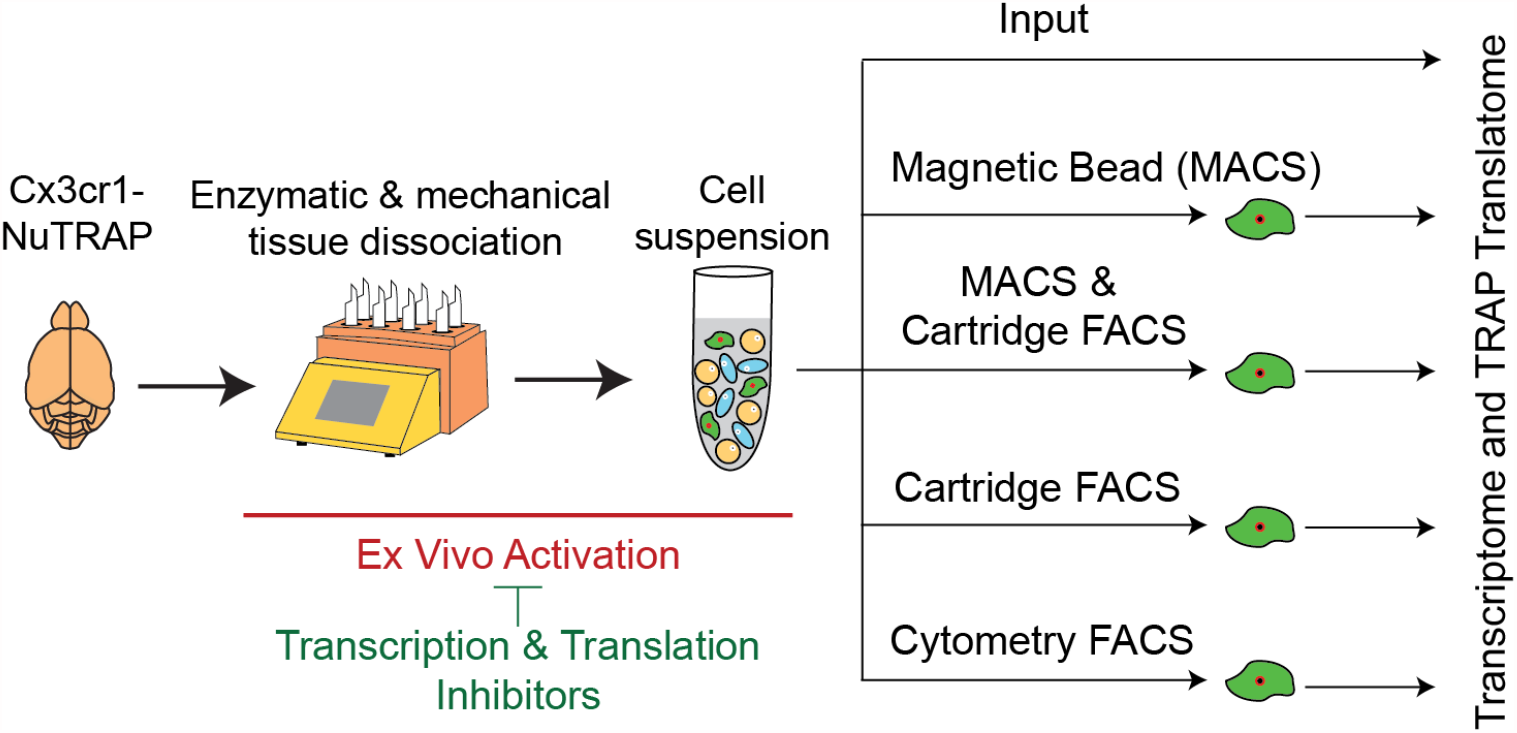

**Main Points:** MACS, cytometer-FACS, and cartridge-FACS give equivalent and sufficient yield/purity for microglial analyses. *Ex vivo* microglial activation is prevented by supplementation with transcription/translation inhibitors during cell preparation.

## Introduction

Microglia, the brain’s resident macrophages, have come to the forefront of neuroimmunology research (Prinz, Jung, & Priller, 2019). They serve as surveyors of the central nervous system and exhibit behavior derived from their embryonic precursors, myeloid cells (Cuadros & Navascues, 1998; Rock et al., 2004), with roles in neurodevelopment, sex differences, as well as in health and neurodegenerative diseases (Butovsky & Weiner, 2018; Han, Fan, Zhou, Blomgren, & Harris, 2021; Salter & Stevens, 2017). Microglial activity is governed by local microenvironments and through communication with neighboring cells. Under stress, microglial cells transition to an activated phenotype, classically defined by morphological transformation from ramified to amoeboid, release of pro-inflammatory cytokines, and a shift in global gene expression (Avignone, Lepleux, Angibaud, & Nagerl, 2015; Rock et al., 2004; Sierra, Abiega, Shahraz, & Neumann, 2013). With the advent of single cell transcriptomic sequencing, the field has undergone a taxonomy reclassification (Dubbelaar, Kracht, Eggen, & Boddeke, 2018; Provenzano, Perez, & Deleidi, 2021). Current evidence suggests microglia exist within a phenotypic gradient, and the transition away from a quiescent state is no longer viewed as binary ‘on’ or ‘off’. Thus, the use of microglial gene expression profiles to infer functional status has bolstered the use of transcriptomic profiling as a powerful technique for microglial classification.

Traditionally, transcriptomic analyses from specific cell types required the liberation of cells from their native environment and use of fluorescence-activated cell sorting (FACS) or MACS labeling techniques prior to RNA extraction (Cahoy et al., 2008). Cell dissociation primarily consists of enzymatic and mechanical dissociation with quality checks for cell viability, debris removal, and absence of cell aggregation (Reichard & Asosingh, 2019). Creation of a single-cell suspension from brain tissue and isolation of microglia is harsh and may alter the phenotypic state of microglia *ex vivo* (Haimon et al., 2018; Wu, Pan, Zuo, Li, & Hong, 2017). *Ex vivo* microglial activation has the potential of introducing confounds that may mask endogenously induced activation, such as in a pathological state. To avoid cell-isolation confounds, microglial-specific cre-lines (Cx3cr1-Cre) have been combined with various floxed ribosomal tagging models: 1) ribosome tagging (RiboTag) (Haimon et al., 2018), 2) translating ribosome affinity purification (TRAP) (Ayata et al., 2018), and 3) nuclear tagging and translating ribosome affinity purification (NuTRAP) (Chucair-Elliott et al., 2020), allowing the immunoprecipitation (IP) of tagged polysomes to isolate microglial-specific translatomes without the need for cell isolation. Although transgenic ribosome IP-approaches overcome many of the potential confounds of *ex vivo* activation, experimental endpoints such as proteomics and single cell heterogeneity still require cell isolation of intact microglial cells. Additionally, animal availability, complex breeding strategies, and cost will continue to be a deterrent for many investigators to using transgenic microglial labeling. While single-cell sequencing allows for broad and potentially unbiased analysis of various cell types, it too is predicated on the creation of a single-cell suspension. Thus, an understanding of the effects of cell preparation and isolation methods on *ex vivo* activation while maintaining highly pure microglial enrichment is needed. The advent of ribosome tagging approaches allows generation of a reference microglial signature to which sorted microglial profiles can be compared. Thus, the goals of this study were to compare purity and yield of isolated microglia and assess the relative level of *ex vivo* activation by comparing Cx3cr1-TRAP-isolated RNA to various sorting techniques used for microglia: MACS (Holt & Olsen, 2016; Nikodemova & Watters, 2012), cytometer-based FACS (Hickman et al., 2013), and newly available low-pressure cartridge-based FACS (Roberts, Anderson, Carmody, & Bosio, 2021)) using TRAP-enrichment as a baseline purified microglial translatome for *ex vivo* activation.

The previously described Cx3cr1-NuTRAP line (Chucair-Elliott et al., 2020) was used in all comparisons of isolation efficacy of various cell sorting techniques including high-and low-pressure fluorescence-activated cell sorting (FACS) as well as magnetic-activated cell sorting (MACS) isolation. We compared artifacts induced through three cell sorting techniques via transcriptomic profiling of bulk tissue, sorted cells, and immunoprecipitated translatomes and found similar performance with *ex vivo* activational signatures principally occurring during the enzymatic digestion and mechanical dissociation during initial cell preparation. Inclusion of transcriptional and translational inhibitors during the cell preparation step prevented most of these artifacts. These studies provide critical insight into the sensitivity of microglia and guidance on experimental design to minimize *ex vivo* confounds of microglial isolation.

## Materials and Methods

### Animals

All animal procedures were approved by the Institutional Animal Care and Use Committee at the Oklahoma Medical Research Foundation (OMRF). Parent mice were purchased from the Jackson Laboratory (Bar Harbor, ME) and bred and housed in the animal facility at the OMRF, under SPF conditions in a HEPA barrier environment. Cx3cr1-Cre/ERT2^+/+^ males (stock # 20940) (Yona et al., 2013) were mated with NuTRAP^flox/flox^ females (stock # 029899) (Chucair-Elliott et al., 2020; Roh et al., 2018) to generate the desired progeny, Cx3cr1-cre/ERT2^+/wt^; NuTRAP^flox/wt^ (Cx3cr1-NuTRAP) (Chucair-Elliott et al., 2020). DNA was extracted from mouse ear punch samples for genotyping. Mice (males and females) were ∼3-8 months old at the time of performing experiments. Euthanasia prior to tissue harvesting was carried out by cervical dislocation followed by rapid decapitation. The primers used for genotyping (Integrated DNA Technologies, Coralville, IA) are included in **Table S1**.

### Tamoxifen (Tam) induction of cre recombinase

At ∼3 months of age, mice received a daily intraperitoneal (ip) injection of tamoxifen (Tam) solubilized in 100% sunflower seed oil by sonication (100 mg/kg body weight, 20 mg/mL stock solution, #T5648; Millipore Sigma, St. Louis, MO) for five consecutive days (Chucair-Elliott et al., 2020; Chucair-Elliott et al., 2019; Srinivasan et al., 2016).

### Preparation of single cell suspension from mouse brain

Halves of Cx3cr1-NuTRAP mouse brains were rinsed in D-PBS, sliced into 4 sagittal sections and placed into gentleMACS C-tubes (#130-093-237, Miltenyi Biotec, San Diego, CA), and processed for generation of single-cell suspensions using the Adult Brain Dissociation Kit and GentleMACS Octo Dissociator system (#130-107-677 and #130-095-937, respectively, Miltenyi Biotec) (Chucair-Elliott et al., 2020). Following debris removal (#130-109-398, Miltenyi Biotec), cells were resuspended in 1 mL 0.5% BSA in D-PBS (#130-091-376, Miltenyi Biotec) and filtered through a 35 µm filter (#352235, Fisher Scientific). An aliquot of cells was retained as “Cell Input” for flow cytometric and RNA-Seq analyses.

### Cell Counting

Filtered cells were diluted 10x in 0.5%BSA in D-PBS (#130-091-376, Miltenyi Biotec) prior to cell counting on a MACSQuant Analyzer 10 (#130-096-343, Miltenyi Biotec). 50 µL diluted cells were analyzed to determine absolute cell count. Cells were gated on FSC-A/SSC-A to determine cell count and FSC-A/FSC-H to determine singlet count. Absolute cell counts were used to determine antibody staining ratios.

### Cd11b Magnetic Labeling and Separation

Cells were pelleted at 300 x g for 10 minutes at 4ºC and resuspended in 90 µL of 0.5% BSA in D-PBS with 10 µL of CD11b (Microglia) MicroBeads (#130-093-636, Miltenyi Biotec) per 10^7^ total cells. After mixing well, cells were incubated for 15 minutes at 2-8°C protected from light. Cells were washed with 1 mL of 0.5% BSA in D-PBS and pelleted at 300 x g for 10 minutes. The cell pellet was resuspended in 500 µL of 0.5% BSA in D-PBS. After priming the autoMACS Pro Separator (#130-092-545, Miltenyi Biotec), sample and collection tubes were placed in a cold Chill5 tube rack (#130-092-951, Miltenyi Biotec) with both positive and negative fractions being collected. The double-positive selection (Posseld) program (ie., positive fraction cells are then run over a second magnetic column for higher purity) was used to elute highly pure Cd11b+ cells in 500 µL. Following separation, the positive fraction was reserved for further applications and analysis.

### Antibody Labeling for FACS

Cell suspensions were pelleted at 300 x g for 10 minutes at 4ºC and resuspended in 96 µL of 0.5% BSA in D-PBS, 2 µL of Cd11b-APC antibody (M1/70, #130-113-793, Miltenyi Biotec), and 2 µL of Cd45-VioBlue antibody (REA737, #130-110-802, Miltenyi Biotec). After mixing well, cells were incubated for 10 minutes in the refrigerator (2-8°C) protected from light. Cells were washed with 1 mL of 0.5% BSA in D-PBS and pelleted at 300 x g for 10 minutes. Cells suspensions from half brains were processed in parallel for Cartridge-Based FACS (MACSQuant Tyto) and Cytometer-Based FACS (FACSAria).

### Cartridge-based FACS (MACSQuant Tyto)

Stained cell pellets were resuspended in 10 mL of 0.5% BSA in D-PBS. A MACSQuant Tyto Cartridge (#130-106-088, Miltenyi Biotec) was primed using 1 mL of 0.5% BSA in D-PBS. The cell suspension was then filtered through 20 µm Pre-Separation Filters (#130-101-812, Miltenyi Biotec). An aliquot of 500 µL of filtered cell suspension was saved as the Tyto Input fraction for analysis. The remaining cell suspension was then loaded into the input chamber of a MACSQuant Tyto cartridge. After loading labeled cells into the input chamber, the cartridge was scanned into the MACSQuant Tyto Cell Sorter system (#130-103-931, Miltenyi Biotec) and sorting parameters were selected. The MACSQuant Tyto cartridge is a sterile, closed, single-use system that relies on accurate activation of the sort valve to pass cells of interest (in this case microglia) into the positive sort chamber. Cell speed (or time-of-flight) was determined by the time it took a cell to pass between two adjacent PMT lasers. In this study, the V1 filter (450/50 nm) of the Violet (405 nm) laser was used as a cell trigger -the first PMT channel used to measure cell speed, to detect Cd45-Vioblue positive cells at a threshold signal of 20. The B1 filter (525/550) of the blue (488 nm) laser was used as the cell speed channel to detect eGFP+ cells at a signal threshold of 4. A blue (488 nm) laser with B1 (525/50 nm) and B2 (585/40 nm) filter combinations was used to gate on eGFP^+^ cells without auto-fluorescence interference. Subsequent gating based on CD11b-APC fluorescence used a red (638 nm) laser and R1 (655-730 nm) filter. The gating strategy was set to Cd11b^+^Cd45^+^eGFP^+^ for isolation of microglia (**Figure S1**). After completion of the sort, the cells from the positive fraction chamber were collected. The positive fraction chamber was washed twice using 450 µL of 0.5% BSA in D-PBS per wash and combined with the initial positive fraction collection. After sorting was completed, an aliquot of (10%) of the positive fraction was kept for analysis on the MACSQuant Analyzer 10 Cytometer.

### Cytometer-based FACS (FACSAria)

Following staining, cells were pelleted and then resuspended in 2 mL of 0.5% BSA in D-PBS for cytometer-based sorting (FACSAria IIIu cell sorter, BD Biosciences). An aliquot of 200 µL stained cells was saved as input for analysis. A Violet (405 nm) laser was used to gate Cd45-Vioblue positive cells using a 450/40 nm filter. A Blue (488 nm) laser with 530/30 nm and Yellow-Green (561 nm) laser with 610/20 nm filter combinations was used to gate on eGFP^+^ cells without auto-fluorescence interference. A Red (640 nm) laser was used to detect Cd11b-APC fluorescence with a 660/20 nm filter. The gating strategy was set to Cd11b^+^Cd45^+^eGFP^+^ for isolation of microglia (**Figure S2**). After sorting was completed, an aliquot (10%) of the positive fraction was kept for analysis on the MACSQuant Analyzer 10 Flow Cytometer.

### Addition of Inhibitors

Transcription and translation inhibitors were included during cell preparation, as previously described (Marsh et al., 2020) with slight modifications. Briefly, Actinomycin D (#A1410, Sigma-Aldrich) was reconstituted in DMSO to a concentration of 5mg/mL before being aliquoted and stored at -20°C protected from light. Triptolide (#T3652, Sigma-Aldrich) was reconstituted in DMSO to a concentration of 10mM before being aliquoted and stored at -20°C protected from light. Anisomycin (#A9789, Sigma-Aldrich) was reconstituted in DMSO to a concentration of 10mg/mL before being aliquoted and stored at 4°C protected from light. For the samples to be treated with inhibitors, 2 µL each of Actinomycin D, Triptolide, and Anisomycin stocks were added to the initial enzyme 1 mixture prior to dissociation.

### Flow Cytometry Analysis

For analysis of cell sorting, aliquots of input and positive fractions from each of the sort methods (AutoMACS, AutoMACS to MACSQuant Tyto, MACSQuant Tyto, FACSAria) were taken for analysis on the MACSQuant Analyzer 10 Flow Cytometer. AutoMACS input and positive fractions were stained with Cd11b-APC (M1/70, #130-113-793, Miltenyi Biotec) and Cd45-Vioblue (REA737, #130-110-802, Miltenyi Biotec) after completion of the sort, according to manufacturer’s instructions. AutoMACS to MACSQuant Tyto, MACSQuant Tyto, and FACSAria input and positive fractions were stained prior to cell sorting. Following staining, cells were resuspended in 500 µL of 0.5% BSA/D-PBS and run on the MACSQuant Analyzer 10 Flow Cytometer. Post-sort purity was assessed by: 1) percent eGFP+ singlets and 2) percent Cd11b+Cd45+ singlets (**Figure S3**) using MACSQuantify v2.13.0 software.

To test the effect of transcription and translation inhibitors on the relative abundance of cell types following cell preparation, aliquots of cells were stained with: 1) Microglial (Cd11b-APC (M1/70, #130-113-793, Miltenyi Biotec) / Cd45-Vioblue (REA737, #130-110-802, Miltenyi Biotec)), 2) Neuronal (Cd24-Vioblue (REA743, #130-110-831, Miltenyi Biotec)), 3) Astrocytic (ACSA2-APC (REA969, #130-116-245, Miltenyi Biotec)), or 4) Oligodendrocytic (O4-APC (REA576, #130-119-982, Miltenyi Biotec)) fluorophore-conjugated antibodies, according to manufacturer’s instructions. Cells were washed and resuspended in 500 µL and run on the MACSQuant Analyzer 10 Flow Cytometer. Relative cell proportions with and without transcription/translation inhibitors were assessed (**Figure S4**) using MACSQuantify v2.13.0 software.

### Translating Ribosome Affinity Purification (TRAP) and RNA extraction

Purification of ribosomal-bound, microglial-specific RNA was performed as previously described (Chucair-Elliott et al., 2020; Kang et al., 2018; Roh et al., 2018) with slight modifications. For TRAP from whole tissue, a hemisected half-brain was minced into small pieces and homogenized in 1.5 mL ice-cold complete homogenization buffer (50 mM Tris, pH 7.4; 12 mM MgCl2; 100 mM KCl; 1% NP-40; 1 mg/mL sodium heparin; 1 mM DTT; 100 µg/mL cycloheximide (#C4859-1ML, Millipore Sigma); 200 units/mL RNaseOUT Recombinant Ribonuclease Inhibitor (#10777019; Thermofisher); 0.5 mM Spermidine (#S2626, Sigma), 1X complete EDTA-free Protease Inhibitor Cocktail (#11836170001; Millipore Sigma)) with a glass dounce tissue grinder set (#D8938; 15 times with pestle A). For TRAP from cells, after pelleting cells at 1000 x g for 10 min at 4°C, cells were resuspended in 400 µl of complete ice-cold homogenization buffer, transferred to a glass dounce tissue grinder set, and homogenized 15 times with pestle A. Volume was brought up to 1.5 mL with complete homogenization buffer. Homogenates (from tissue or cells) were transferred to 2 mL round-bottom tubes and centrifuged at 12,000 x g for 10 minutes at 4°C. After centrifugation, 100 µL of the supernatant was saved as “RNA” input. The remaining supernatant was transferred to a 2 mL round-bottom tube and incubated with 5 µg/µl of anti-GFP antibody (ab290; Abcam) at 4°C with end-over-end rotation for one hour. Dynabeads Protein G for Immunoprecipitation (#10003D; Thermofisher) were washed three times in 1 mL ice-cold low-salt wash buffer (50mM Tris, pH 7.5; 12mM MgCl2; 100mM KCl; 1% NP-40; 100μg/mL cycloheximide; 1mM DTT). After removal of the last wash, the homogenate/antibody mixture was transferred to the 2 mL round-bottom tube containing the washed Protein-G Dynabeads and incubated at 4°C with end-over-end rotation overnight. Magnetic beads were collected using a DynaMag-2 magnet and the unbound-ribosomes and associated RNA discarded. Beads and GFP-bound polysomes were then washed three times with 0.5 mL of high-salt wash buffer (50mM Tris, pH 7.5; 12mM MgCl2; 300mM KCl; 1% NP-40; 100μg/mL cycloheximide; 2mM DTT). Following the last wash, 350 µL of Buffer RLT (Qiagen, Germantown, MD) supplemented with 3.5 µl 2-β mercaptoethanol (#444203, Sigma) was added directly to the beads and incubated with mixing on a ThermoMixer (Eppendorf) for 10 minutes at room temperature. The beads were magnetically separated and the supernatant containing the target bead-bound polysomes and associated RNA was transferred to a new tube. 350 µl of 100% ethanol was added to the tube (“TRAP” fraction: enriched in transcriptome associated to EGFP-tagged ribosomes) and then loaded onto a RNeasy MinElute column (Qiagen). RNA was isolated using RNeasy Mini Kit (#74104, Qiagen), according to manufacturer’s instructions. RNA was quantified with a Nanodrop 2000c spectrophotometer (Thermofisher Scientific) and its quality assessed by HSRNA screentape with a 2200 Tapestation analyzer (Agilent Technologies).

### Library construction and RNA sequencing (RNA-seq)

Directional RNA-Seq libraries were made from 5-100 ng RNA, as previously described (Chucair-Elliott et al., 2020; Chucair-Elliott et al., 2019). Briefly, poly-adenylated RNA was captured using NEBNext Poly(A) mRNA Magnetic Isolation Module (#NEBE7490L; New England Biolabs, Ipswich, MA) and then processed using NEBNext Ultra II Directional Library Prep Kit for Illumina (#NEBE7760L; New England Biolabs) for the creation of cDNA libraries, according to the manufacturer’s instruction. Library sizing was performed with HSRNA ScreenTape (#5067-5579; Agilent Technologies) and libraries were quantified by Qubit HSDNA (#Q32851, Thermo). The libraries for each sample were pooled at 4 nM concentration and sequenced using an Illumina NovaSeq 6000 system (SP PE50bp, S4 PE150) at the OMRF Clinical Genomics Center. The entirety of the sequencing data is available for download in FASTQ format from NCBI Sequence Read Archive (GSE179721).

### RNA-Seq Data Analysis

Following sequencing, reads were trimmed and aligned prior to differential expression statistics and correlation analyses in Strand NGS software package (v4.0) (Strand Life Sciences). Reads were aligned against the mm10 build of the mouse genome (2014.11.26). Alignment and filtering criteria included: adapter trimming, fixed 2bp trim from 5’ and 2bp from 3’ ends, a maximum number of one novel splice allowed per read, a minimum of 90% identity with the reference sequence, a maximum of 5% gap, and trimming of 3’ end with Q<30. Alignment was performed directionally with Read 1 aligned in reverse and Read 2 in forward orientation. Normalization was performed with the DESeq2 algorithm (Love, Huber, & Anders, 2014). Transcripts with an average read count value >5 in at least 100% of the samples in at least one group were considered expressed at a level sufficient for quantitation per tissue and those transcripts below this level were considered not detected/not expressed and excluded, as these low levels of reads are close to background and are highly variable. For statistical analysis of differential expression, a one-way ANOVA or two-way ANOVA with the factors of TRAP fraction and treatment and a Benjamini-Hochberg Multiple Testing Correction followed by Student-Newman Keuls post hoc test were used (FDR<0.1). For those transcripts meeting this statistical criterion, a fold change >|2| cutoff was used to eliminate those genes which were statistically significant but unlikely to be biologically significant and orthogonally confirmable due to their very small magnitude of change. Visualizations of hierarchical clustering and principal component analyses were performed in Strand NGS (Version 3.1, Bangladore, India). Cell type specific marker gene lists were generated from the re-analysis published by McKenzie et al. (McKenzie et al., 2018) of immunopurified and high throughput single cell data from mice as we have described previously (Chucair-Elliott et al., 2020). Over-representation analysis (ORA) was conducted using WEB-based Gene SeT AnaLysis Toolkit (WebGestalt, www.webgestalt.org)(Liao, Wang, Jaehnig, Shi, & Zhang, 2019; Wang, Duncan, Shi, & Zhang, 2013; Wang, Vasaikar, Shi, Greer, & Zhang, 2017; B. Zhang, Kirov, & Snoddy, 2005). Top over-represented biological processes were selected from gene ontology functional database with no redundant option selected (Hypergeometric test, BHMTC, FDR<0.05) and background reference gene list of all expressed genes (raw count>5 in all samples from at least one group). Top over-represented transcription factor targets were selected from network functional database with all expressed genes as the reference gene list (Hypergeometric test, BHMTC, FDR<0.05). Heatmaps of over-represented biological processes were created using Mopheus (https://software.broadinstitute.org/morpheus). Upset plot was created using UpSetR v 1.4.0 package (Conway, Lex, & Gehlenborg, 2017) in RStudio v 1.4.1106 with R v 4.0.5. Previously published microglial *ex vivo* activational lists were compared (Ayata et al., 2018; Haimon et al., 2018; Marsh et al., 2020) and genes included in at least two of the three previous studies were considered “*ex vivo* activational transcripts”.

### Immunochemistry and imaging

Brain samples were fixed for 4h in 4% PFA, cryoprotected by sequential incubations in PBS containing 15% and 30% sucrose, and then frozen in Optimal Cutting Temperature medium (#4583, Tissue-Tek). Twelve µm-thick sagittal sections were cryotome-cut (Cryostar NX70, ThermoFisher Scientific). Tissue sections were rinsed with PBS containing 1% Triton X-100, blocked for 1h in PBS containing 10% normal donkey serum, and processed for fluorescence immunostaining and downstream analysis, as previously described (Chucair-Elliott et al., 2020). The primary antibodies included rabbit anti-GFP (#ab290, 1:100, Abcam, Cambridge, MA), rat anti-CD11b (#C227, Clone M1/70, 1:100, Leinco Technologies, St. Louis, MO), and rat anti-CD45 (#550539, Clone 30-F111, 1:100, BD Biosciences). Sequential imaging was performed on an Olympus FluoView confocal laser-scanning microscope (FV1200; Olympus; Center Valley, PA) at the Dean McGee Eye Institute imaging core facility at OUHSC. Microscope and FLUOVIEW FV1000 Ver. 1.2.6.0 software (Olympus) settings were identical for samples using the same staining-antibody combination and at same magnification. The experimental format files were oib. The Z-stack generated was achieved at 1.26 µm step size with a total of 8 optical slices at 20X magnification (2X zoom).

## Results

The goal of this study was to compare microglial sorting techniques and determine the relative levels of *ex vivo* activation induced during cell preparation and microglial isolation. A schematic of the experimental design is represented in **Figure 1A**. In the Cx3cr1-NuTRAP mice, following cre recombination in Cx3cr1+ cells, deletion of the floxed stop cassette causes activation of the NuTRAP allele, labeling microglial ribosomes with eGFP and nuclei with biotin and mCherry (Chucair-Elliott et al., 2020). For the first part of the present study, we used eGFP as a sorting criterion and in the evaluation of post-sort microglial purity, along with Cd11b and Cd45 co-expression. Colocalization of eGFP with microglial markers Cd11b and Cd45 in Cx3cr1-NuTRAP brains was verified by immunohistochemistry (**Figure S5**). Enzymatic and mechanical dissociation of Cx3cr1-NuTRAP brains was performed to generate single-cell suspensions.

**Figure 1.**
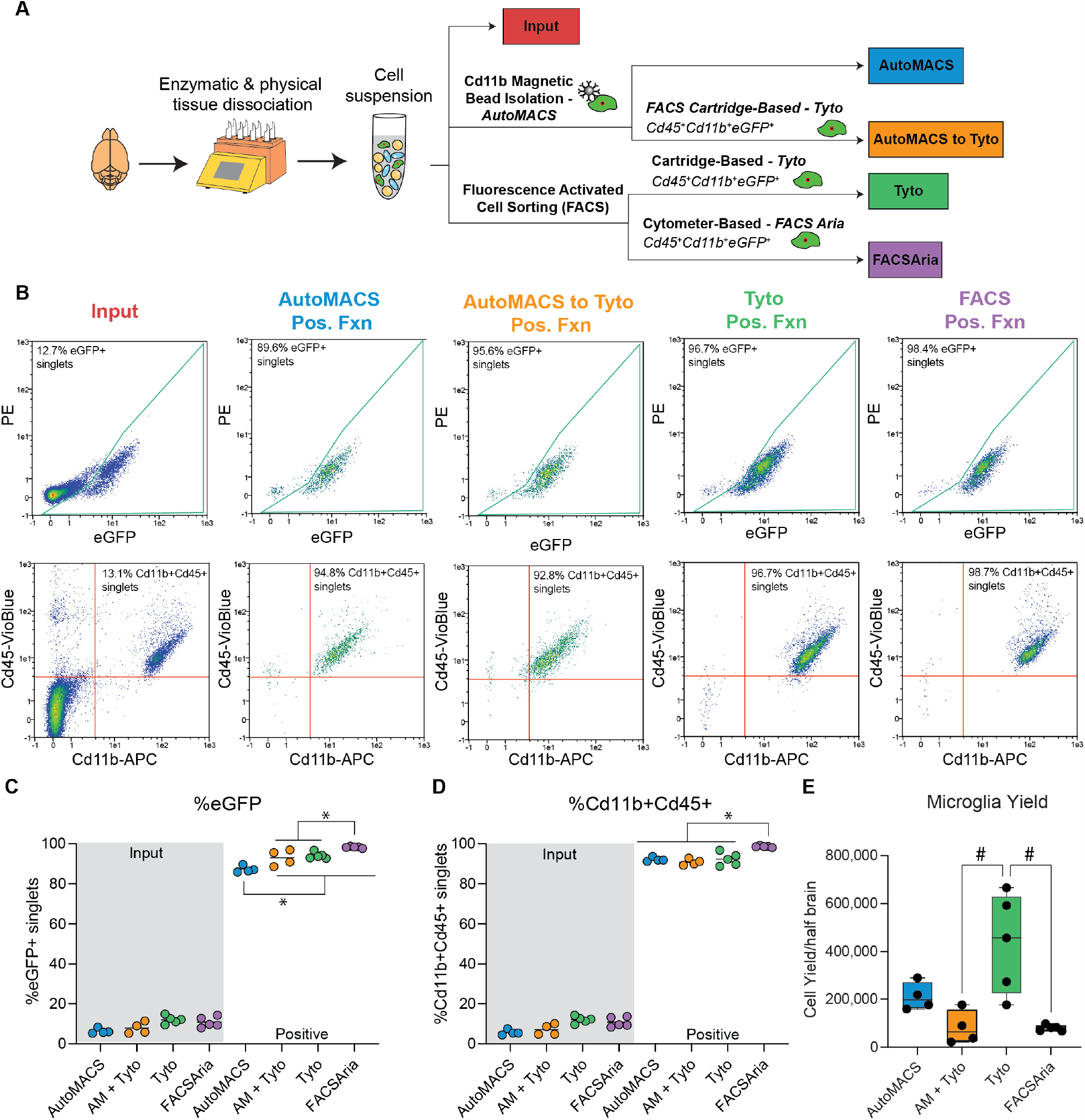
Comparison of purity and yield among microglial cell isolation techniques. A) Schematic of the experimental design. Cx3cr1-NuTRAP brains were enzymatically and mechanically dissociated to create a single cell suspension. Different microglial sorting techniques were compared to cell input for purity, yield, and transcriptomic signatures. B) Representative flow cytometry plots of immunostained single-cell suspensions from input and after each of the sorting strategies shows a distinct population of: eGFP+ cells (identified as Cx3cr1+ microglia) and Cd11b+Cd45+ cells (identified as microglia per traditional cell surface markers). All sort positive fractions were enriched for: (C) eGFP+ singlets and (D) Cd11b+Cd45+ singlets in the positive fractions (as compared to input). (Two-Way ANOVA, Main effect of TRAP Fraction, p<0.001). When comparing positive fractions, the AutoMACS positive fraction had lower %eGFP+ singlets as compared to all other sort methods. FACSAria had higher percentage of eGFP+ singlets than all other sort methods. FACSAria had higher percentage of Cd11b+Cd45+ singlets as compared to all other sort methods positive fractions (Two-way ANOVA, Tukey’s post-hoc, *p<0.05). E) Microglial yield was significantly higher in the MACSQuant Tyto positive fraction as compared to the AutoMACS to MACSQuant Tyto and FACSAria positive fractions (One-Way ANOVA, Tukey’s posthoc, #p<0.01).

### Flow cytometric analysis of sort fractions from various microglial sorting techniques

After reserving an aliquot as input, cells were subjected to one of four isolation techniques: 1) Cd11b+ magnetic-bead based isolation (AutoMACS), 2) Cartridge-based FACS on Cd11b+/Cd45+/eGFP+ (MACSQuant Tyto), 3) Cytometer-based FACS on Cd11b+/Cd45+/eGFP+ (FACSAria), and 4) AutoMACS debulking of cells prior to cartridge-based FACS (AutoMACS to MACSQuant Tyto) (**Figure 1A**).

Aliquots of cell input and positive sort fractions from each of the four sort methods were analyzed by flow cytometry. All sort methods showed enriched populations of eGFP+ and Cd11b+/Cd45+ singlets in their positive fractions as compared to cell input (**Figure 1B**). The positive fractions of all sort methods were enriched in eGFP+ singlets as compared to the input fraction (**Figure 1C**; Two-way ANOVA, main effect of sort fraction, ***p<0.001). The AutoMACS sort resulted in lower overall percentage of eGFP+ singlets compared to all other sort methods and the FACSAria sort resulted in a higher overall percentage of eGFP+ singlets, though all approaches demonstrated a high level of enrichment (**Figure 1C;** Two-way ANOVA, Tukey’s post-hoc, *p<0.05). The positive fractions of all sort methods were enriched in Cd11b+Cd45+ singlets as compared to the input fraction (**Figure 1D**; Two-way ANOVA, Main effect of sort fraction, p<0.001). FACSAria sort resulted the highest overall percentage of Cd11b+Cd45+ singlets (**Figure 1D;** Two-way ANOVA, Tukey’s post-hoc, *p<0.05). Although FACSAria had higher microglial purity than other sort methods, it showed a significantly lower yield than the MACSQuant Tyto sort (**Figure 1E**; One-way ANOVA, Tukey’s post-hoc, #p<0.05).

### Comparison of transcriptomic profiles of microglia isolated from various sort methods

Following cell preparation and isolation using methods displayed in Figure 1A, RNA was isolated from cells for preparation of stranded RNA-Seq libraries. We first examined enrichment/depletion of previously published microglial, astrocytic, oligodendrocytic, neuronal, and endothelial markers in the transcriptomic profiles (Chucair-Elliott et al., 2020; McKenzie et al., 2018) (**Supplemental Table 1**). Each of the four sort methods showed similar levels of enrichment of microglial marker genes (**Figure 2A**) and depletion of astrocytic, oligodendrocytic, neuronal, and endothelial marker genes (**Figure 2B-E**) when compared to cell input. In combination with the flow cytometric data presented in Figure 1, this gives confidence that each of the sort methods are effective in isolating highly pure populations of microglia.

**Figure 2.**
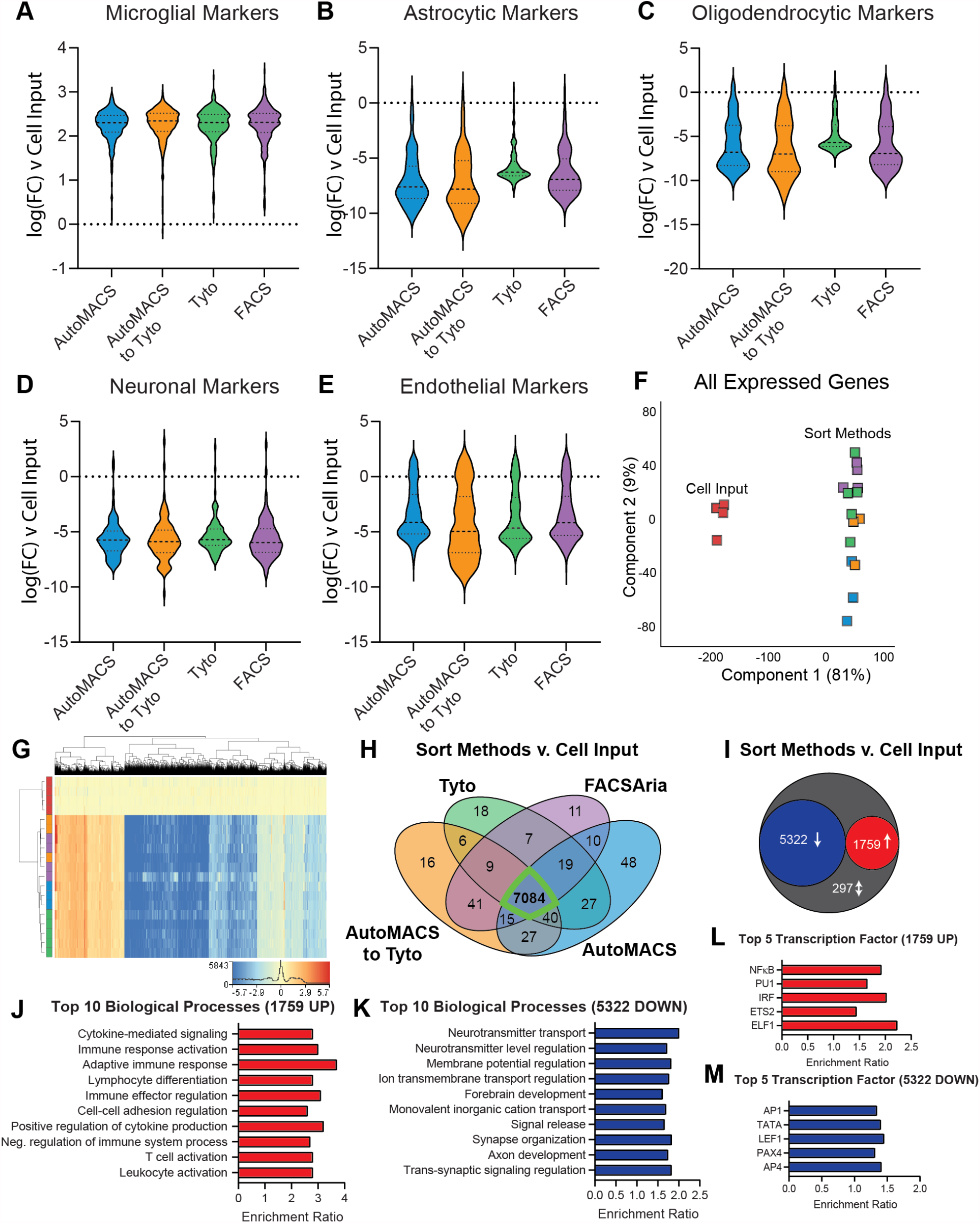
Comparison of transcriptomic profiles of microglia isolated from various cell sorting methods. RNA-Seq libraries were made from each of the groups represented in Figure 1A to compare the transcriptomic profiles of microglia isolated via four different cell sorting strategies. Each of the strategies had similar levels of (A) enrichment of microglial transcripts and depletion of: (B) astrocytic, (C) oligodendrocytic, (D) neuronal, and (E) endothelial transcripts when compared to cell input. F) Principal component analysis of all expressed genes shows clear separation of cell input from all other sort methods in the first component with 81% of explained variance. G) Hierarchical clustering of differentially expressed genes (DEGs) (One-Way ANOVA, BHMTC, SNK FDR<0.1, |FC|>2) shows separation of cell input and sort methods. Each of the sort methods show very similar patterning of expression across DEGs. H) Comparison of SNK post-hocs from each of the sort methods v. cell input, showed the majority of enrichments/depletions (ie.,DEGs) (7084/7378 = 96%) were in common between all sort methods. I) There were 5322 DEGs (72%) that were depleted and 1759 DEGs (24%) that were enriched in all sort methods compared to cell input. There were only 297 discordant DEGs (4%) between the different sort methods as compared to cell input. J) Top 10 biological processes over-represented in the 1759 genes upregulated in all sort methods compared to cell input (Gene Ontology Over-Representation Analysis, BHMTC FDR <0.05). K) Top 10 biological processes over-represented in the 5322 genes downregulated in all sort methods compared to cell input (Gene Ontology Over-Representation Analysis, BHMTC FDR <0.05). L) Top 5 transcription factor targets over-represented in the 1759 genes upregulated in all sort methods compared to cell input (Transcription factor target network over-representation analysis, BHMTC FDR<0.05). M) Top 5 transcription factor targets over-represented in the 5322 genes downregulated in all sort methods compared to cell input (Transcription factor target network over-representation analysis, BHMTC FDR<0.05).

Next, we examined the transcriptomic data in an unbiased manner. Principal component analysis of all expressed genes (>5 counts in all samples from at least one group) showed clear separation of cell input from all sort methods in the first component with 81% of the explained variance (**Figure 2F**). Differentially expressed genes were called by One-way ANOVA with Benjamini-Hochberg multiple testing correction (BHMTC) followed by Student-Newman Keuls post hoc (FDR<0.1, |FC|>2; **Supplemental Table 2**). Hierarchical clustering of the 7378 DEGs shows separation of cell input from all sort methods with similar patterning of enrichment and depletion of DEGs across all sort methods scaled to cell input (**Figure 2G**). The majority of pairwise DEGs (sort method v cell input) were in common between all sort methods (7084/7378 = 96%) (**Figure 2H**), suggesting a high degree of similarity between each of the sort methods. In addition, 5322 DEGs (72%) that were down and 1759 DEGs (24%) were up in all sort methods compared to cell input. There were 297 discordant DEGs (4%) between the different sort methods as compared to cell input (**Figure 2I**).

Over-representation analysis of gene ontology (ORA GO) of the 1759 genes that were up across all sort methods identified 177 over-represented biological processes pathways (BHMTC, FDR<0.05; **Supplemental Table 3**). Examination of the top 10 over-represented biological processes reveals several pathways involved in microglial function, including: cytokine-mediated signaling, immune response activation, and adaptive immune response, among others (**Figure 2J**). Running a similar ORA GO on the 5322 genes down across all sort methods (compared to cell input) revealed 252 over-represented biological processes (BHMTC, FDR<0.05; **Supplemental Table 3**). The top 10 processes include many neuron-focused pathways, such as: neurotransmitter transport, neurotransmitter level regulation, and membrane potential regulation (**Figure 2K**), indicating depletion of these genes in the sorted cells.

Next, we examined the common 1759 up-regulated and 5322 down-regulated genes across the four sort methods for over-representation of transcription factor targets. Network ORA on the 1759 genes enriched in the positive fraction of all sort methods identified 21 over-represented transcription factor targets, including the top five hits: Elf1, Ets2, Irf, Pu1, and Nfkb (**Figure 2L, Supplemental Table 3**). Three of the top five transcription factor targets (Elf1, Ets2, PU.1) are part of the ETS family of transcription factors that assist in regulating immunity (Gallant & Gilkeson, 2006), with PU.1 being a “master regulator” of microglial identity and function (Yeh & Ikezu, 2019). The other two transcription factors (Irf and Nfkb) are also critical regulators of inflammation and antiviral response (Iwanaszko & Kimmel, 2015), an important function of microglia.

Network ORA of the 5322 genes down in all sort methods compared to input identified 468 over-represented transcription factor targets, including top five hits: Ap1, Tata, Lef1, Pax4, and Ap4. Ap1 transcription factors are of the Jun and Fos family and have been shown to interact with Brain-derived neurotrophic factor (BDNF) to modulate neuronal synaptic plasticity. Lef1 is an endothelium-specific transcription factor (Hupe et al., 2017). Ap4 is an adaptor protein complex that is involved in vesicular trafficking of membrane proteins. Lack of Ap4 has been shown to cause accumulation of axonal autophagosomes containing AMPA receptor components in hippocampal neurons and cerebellar Purkinje cells (Matsuda et al., 2008). Overall, the top transcription factor targets of the genes depleted in the sort fractions (compared to input) are non-microglial regulators.

In combination, the flow cytometric data, distribution of marker gene enrichments/depletion, and analysis of differentially expressed genes (including pathway and transcription factor analysis) suggest that each of the sort methods are producing highly pure populations of microglia with very similar transcriptomic profiles.

### Comparison of TRAP-isolated microglial translatome from tissue homogenate, cell suspension, and various microglial sort methods

A schematic of the experimental design is represented in **Figure 3A**. Cx3cr1-NuTRAP brains were hemisected and processed in halves for whole-tissue homogenization or enzymatic and mechanical dissociation to create single cell suspensions. Single cell suspensions were then sorted using one of four methods: 1) Cd11b magnetic-bead based isolation (AutoMACS), 2) Cartridge-based FACS on Cd11b+/Cd45+/eGFP+ (MACSQuant Tyto), 3) Cytometer-based FACS on Cd11b+/Cd45+/eGFP+ (FACSAria), or 4) AutoMACS debulking of cells prior to cartridge-based FACS (AutoMACS to MACSQuant Tyto), as before. Tissue homogenate (Tissue-TRAP), mixed-cell suspension (Cell-TRAP), and sorted microglia (Sort-TRAP) were then subjected to TRAP pull-down of microglial-specific translating RNA for creation of RNA-Seq libraries.

**Figure 3.**
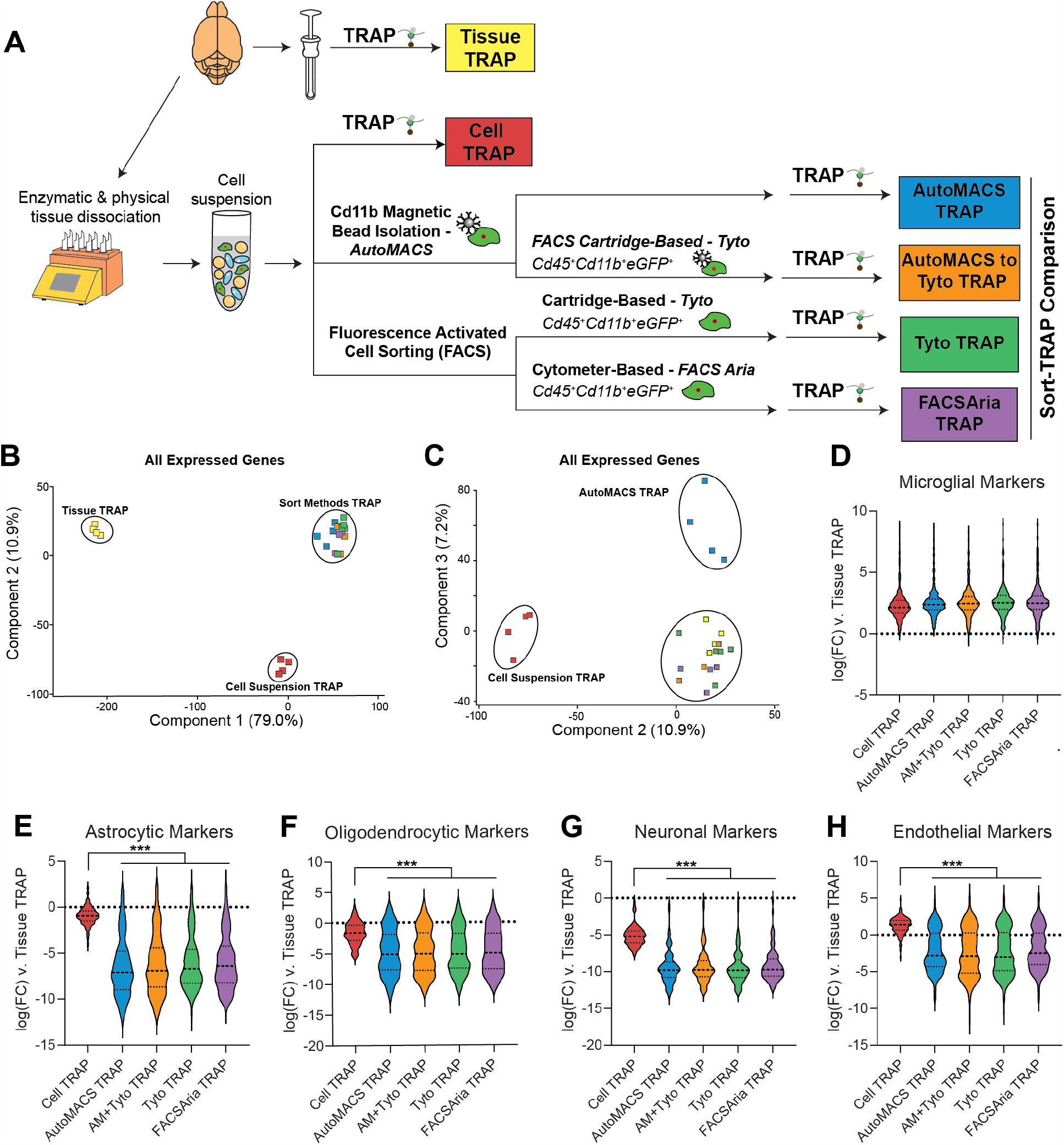
Comparison of TRAP-isolated microglial translatome from tissue homogenate, cell suspension, and various microglial cell sorting methods. A) Schematic of the experimental design. Cx3cr1-NuTRAP brains were hemisected and processed in halves for whole-tissue homogenization or enzymatic and mechanical dissociation to create a single cell suspension. Single cell suspensions were then sorted using MACS-and/or FACS-based isolation of microglia. Tissue homogenate, mixed-cell suspension, and microglia sorted by each of the four depicted methods were subjected to TRAP to isolate microglial-specific ribosomally-bound RNA for creation of RNA-Seq libraries. B) PCA of all expressed genes (>5 read counts in all samples from at least one group) separates Tissue TRAP from all other groups in the first component (79% explained variance) and Cell Suspension TRAP from all other groups in the second component (10.9% explained variance). C) Third component of PCA on all expressed genes separated AutoMACS TRAP from all other groups (7.2% explained variance). Each of the sort strategies had similar levels of (D) enrichment of microglial transcripts and depletion of: (E) astrocytic, (F) oligodendrocytic, (G) neuronal, and (G) endothelial transcripts when compared to Tissue TRAP. All of the sort methods showed stronger depletion of: (E) astrocytic, (F) oligodendrocytic, (G) neuronal, and (G) endothelial transcripts when compared to Cell TRAP (One-way ANOVA, Tukey’s post-hoc, ***p<0.001).

Each of the four Sort-TRAP methods showed similar levels of enrichment of microglial marker genes (**Figure 3D**). Depletion of astrocytic, oligodendrocytic, neuronal, and endothelial marker genes (**Figure 3E-H**) was greater in Sort-TRAP methods as compared to Cell-TRAP (One-way ANOVA, Tukey’s post-hoc, ***p<0.001). This shows that the extra enrichment step of sorting microglia followed by TRAP-isolation of translating microglial RNA, leads to more pure microglial RNA than Tissue-TRAP or Cell-TRAP alone.

In recent years, several studies have suggested that cell-isolation methods cause *ex vivo* activational effects in microglia (Ayata et al., 2018; Haimon et al., 2018; Marsh et al., 2020). In this section, our goal was to determine if different sorting techniques result in different levels of *ex vivo* activation. We used Tissue-TRAP as an “unactivated” microglial reference group, since the Tissue-TRAP method does not rely on the creation of a cell suspension or cell sorting techniques. There were 8076 DEGs between the translatomes when Cell-and Sort-TRAP methods were compared to the Tissue-TRAP reference (One-way ANOVA, BHMTC, SNK FDR<0.1, |FC|>2). Upset plot of the 8076 DEGs shows the majority of the DEGs (7800/8076=97%) are in common between all groups (Cell-/Sort-TRAP v. Tissue-TRAP) (**Figure 4A, Supplemental Table 5**). Hierarchical clustering of all 8076 DEGs shows distinct clustering of Tissue-TRAP from all other groups. Cell-TRAP also clusters separately from the Sort-TRAP groups. These data, together with cell-type enrichments from **Figure 3D-H**, suggests that the act of creating a cell-suspension is the largest contributor to differences seen between Tissue-TRAP and Sort-TRAP methods.

**Figure 4.**
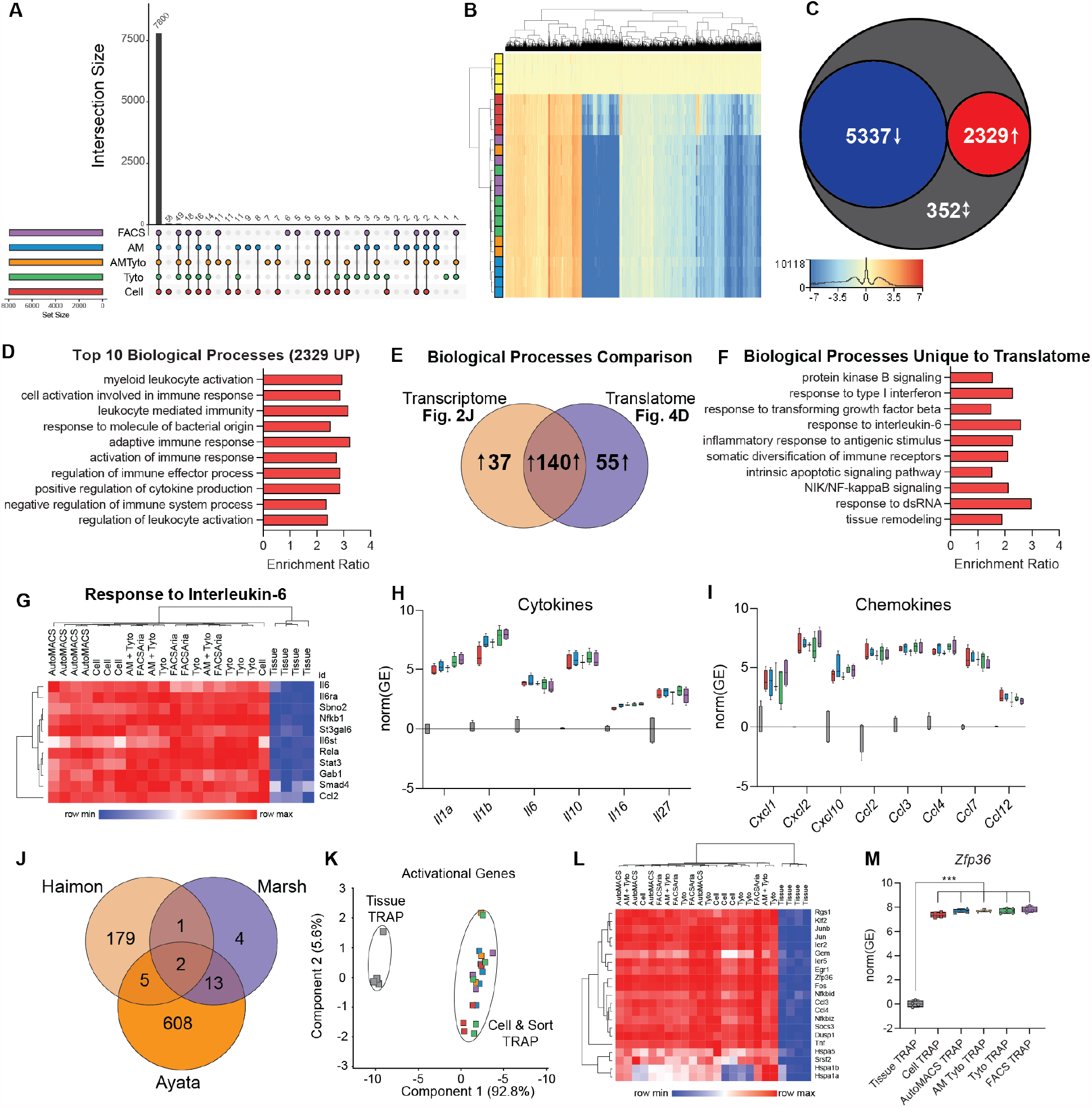
Cell isolation and sorting techniques alter TRAP-isolated microglial translatome compared to whole-tissue TRAP. RNA-Seq libraries were made from each of the groups represented in Figure 3A to compare the TRAP-isolated microglial translatomes between whole-tissue-TRAP and each of the cell isolation and sorting methods. A) Upset plot of DEGs for all groups v. Tissue TRAP (One-Way ANOVA, BHMTC, SNK FDR<0.1, |FC|>2). B) Hierarchical clustering of DEGs shows separation of tissue TRAP from all other groups. Cell TRAP also clusters separately from all Sort-TRAP groups. C) Comparison of DEGs from each group (Cell-and Sort-TRAP) v. Tissue-TRAP, revealed 5337 common DEGs (67%) that were depleted and 2329 common DEGs (29%) that were enriched in all groups (Cell-and Sort-TRAP) compared to Tissue-TRAP. There were only 352 discordant DEGs (4%) between the different sort methods as compared to cell input. D) Top 10 biological processes over-represented in the 2329 genes upregulated in Cell TRAP and Sort-TRAP compared to Tissue TRAP (Gene Ontology Over-Representation Analysis, Hypergeometric test, BHMTC FDR <0.05). E) Comparison of upregulated transcriptomic pathways (Figure 2J; Supplemental Table 3) and upregulated translatome pathways (Figure 4D, Supplemntal Table 5) reveal 55 biological processes that are upregulated in the translatome only. F) Selection of 10 biological processes that are uniquely upregulated in the translatome (from the 55 identified in Figure 4E). G) Heatmap of genes involved in “Response to Interleukin-6 (GO:0070741)” biological process. H) Cytokines (*Il1a, Il1b, Il6, Il10, Il16, Il27*) that are upregulated in the Cell-and Sort-TRAP groups compared to Tissue-TRAP (One-Way ANOVA, BHMTC, SNK FDR<0.1, |FC|>2). I) Chemokines (*Cxcl1, Cxcl2, Cxcl10, Ccl2, Ccl3, Ccl4, Ccl7, Ccl12*) that are upregulated in the Cell-and Sort-TRAP groups compared to Tissue-TRAP (One-Way ANOVA, BHMTC, SNK FDR<0.1, |FC|>2). J) Intersection of activational genes identified in three previous studies (Ayata et al., 2018; Haimon et al., 2018; Marsh et al., 2020) identified 21 *ex vivo* activational transcripts represented in at least two of the studies. I) PCA of 21 *ex vivo* activational genes shows clear separation of tissue TRAP from all other groups in the first component (92.8% explained variance). J) Heatmap of 21 activational genes shows high levels of *ex vivo* activational transcripts across Cell-and Sort-TRAP methods compared to Tissue-TRAP. K) *Zfp36* is enriched in Cell TRAP and Sort-TRAP compared to Tissue TRAP (One-Way ANOVA, Tukey’s posthoc, ***p<0.001).

Next, we identified over-represented pathways among the up-regulated genes compared to Tissue-TRAP. The top 10 biological processes of the 2329 genes up-regulated in comparison to Tissue-TRAP were microglial-related pathways, including: myeloid leukocyte activation, cell activation involved in immune response, and leukocyte-mediated immunity (**Figure 4D**; **Supplemental Table 5**). Comparing the biological processes identified in the transcriptomic analysis (**Figure 2J**; **Supplemental Table 2**) and the translatomic analysis (**Figure 4D**; **Supplemental Table 5**) revealed 55 biological processes that were only up-regulated in the translatome (**Figure 4E**). Several of the biological processes uniquely upregulated in the translatome comparisons were involved in microglial activation pathways, including: response to type I interferon (IFN), response to transforming growth factor beta (TGFB), response to interleukin-6 (IL-6), and NIK/NF-kappaB signaling (NFKB) (**Figure 4F**). IL-6 is a pro-inflammatory cytokine extensively studied in brain aging and disease (Borovcanin et al., 2017; Singh-Manoux et al., 2014; Ye & Johnson, 1999). Microglia have higher Il-6 receptor (Il-6R) expression than any other cell type. As such, microglia are highly responsive to IL-6 and transition into a “primed” state when exposed to high levels of IL-6 (Garner, Amin, Johnson, Scarlett, & Burton, 2018). Hierarchical clustering of the “Response to IL-6” pathway genes showed overall higher levels of expression in Cell-and Sort-TRAP groups (**Figure 4G**; **Supplemental Table 5**). The Cell-TRAP did not cluster separately from the Sort-TRAP groups, providing further evidence that the *ex vivo* activational signature is a function of cell preparation.

Next, we looked at gene expression of common cytokines (**Figure 4H**; **Supplemental Table 5**) and chemokines (**Figure 4I**; **Supplemental Table 5**) across Tissue-, Cell-, and Sort-TRAP groups. We observed higher levels of cytokine and chemokine transcripts across all Cell-and Sort-TRAP groups when compared to Tissue-TRAP (One-way ANOVA, BHMTC, SNK FDR<0.1,

|FC|>2). In an effort to cross-validate our finding with previous studies, we intersected *ex vivo* microglial activational gene lists from three previous studies (Ayata et al., 2018; Haimon et al., 2018; Marsh et al., 2020) and identified 21 *ex vivo* activational transcripts represented in at least two of the studies (**Figure 4J**; **Supplemental Table 5**). PCA of the TRAP data from the present study on the 21 *ex vivo* activational genes shows clear separation of Tissue-TRAP from all other groups in the first component (92.8% explained variance) (**Figure 4K**). Again, suggesting that the *ex vivo* signature is a function of cell preparation. Hierarchical clustering of the 21 activational genes, shows similar patterning as in the “Response to IL-6” pathway with higher levels of expression across Cell-and Sort-TRAP groups compared to Tissue-TRAP (**Figure 4L**; **Supplemental Table 5**). *Zfp36* was one of the *ex vivo* activational genes identified in all three previous studies (Ayata et al., 2018; Haimon et al., 2018; Marsh et al., 2020). Zinc finger protein 36 (*Zfp36*) encodes for the protein Tristetraprolin (TTP) which is involved in regulating immune responses through mRNA destabilizing and alternative splicing (Tu et al., 2019). Zfp36 is enriched in Cell-and Sort-TRAP compared to Tissue TRAP (One-Way ANOVA, Tukey’s posthoc, ***p<0.001).

In summary, these data suggest that *ex vivo* microglial activation is primarily occurring during cell preparation and is sustained through microglial isolation by various sort methods but there were no major differences between the different sort methods.

### Changes in cellularity and *ex vivo* activational profiles following cell preparation

Since enzymatic and mechanical dissociation during cell preparation induced *ex vivo* activational artifacts, we next compared the cellularity and *ex vivo* activational profiles using whole-tissue homogenization and enzymatical/mechanical cell preparation (**Figure 5A**). Normally, flow cytometry is the method of choice in estimating relative cell proportions within a cell suspension. However, these proportions do not account for biased cellular loss during cell preparation. CIBERSORTx, or digital cytometry, estimates cell-type abundance from bulk transcriptomics. Using publicly available cell-type specific data from mouse brain (Y. Zhang et al., 2014) (GSE52564) as a digital cytometry reference matrix, we estimated cell proportions in Tissue Input and Cell Input. There was a decrease in estimated neuronal abundance between Tissue Input (86.4%) and Cell Input (8.7%) and an increase in estimated microglial, astrocytic, endothelial, and oligodendrocytic abundance with cell preparation (**Figure 5B**; **Supplemental Table 6**). Consistent with our CIBERSORTx results, Cell Input was enriched for microglial, astrocytic, oligodendrocytic, and endothelial cell transcripts compared to Tissue Input. Cell-TRAP revealed greater enrichment of microglial cell markers and depletion of astrocytic and oligodendrocytic cell markers than in Tissue-TRAP. (**Figure 5C**; **Supplemental Table 6**) (One-way ANOVA, Tukey’s post-hoc, **p<0.01, **p<0.001). Examining *ex vivo* activational signature genes (Cytokines, Chemokines, Response to IL-6) reveals strong induction with cell preparation (**Figure 5D**; **Supplemental Table 6**). The 21-common *ex vivo* activation signature genes also showed a similar pattern of expression with cell preparation, with high expression in Cell-Input and Cell-TRAP as compared to their tissue counterparts (**Figure 5E**; **Supplemental Table 6**).

**Figure 5.**
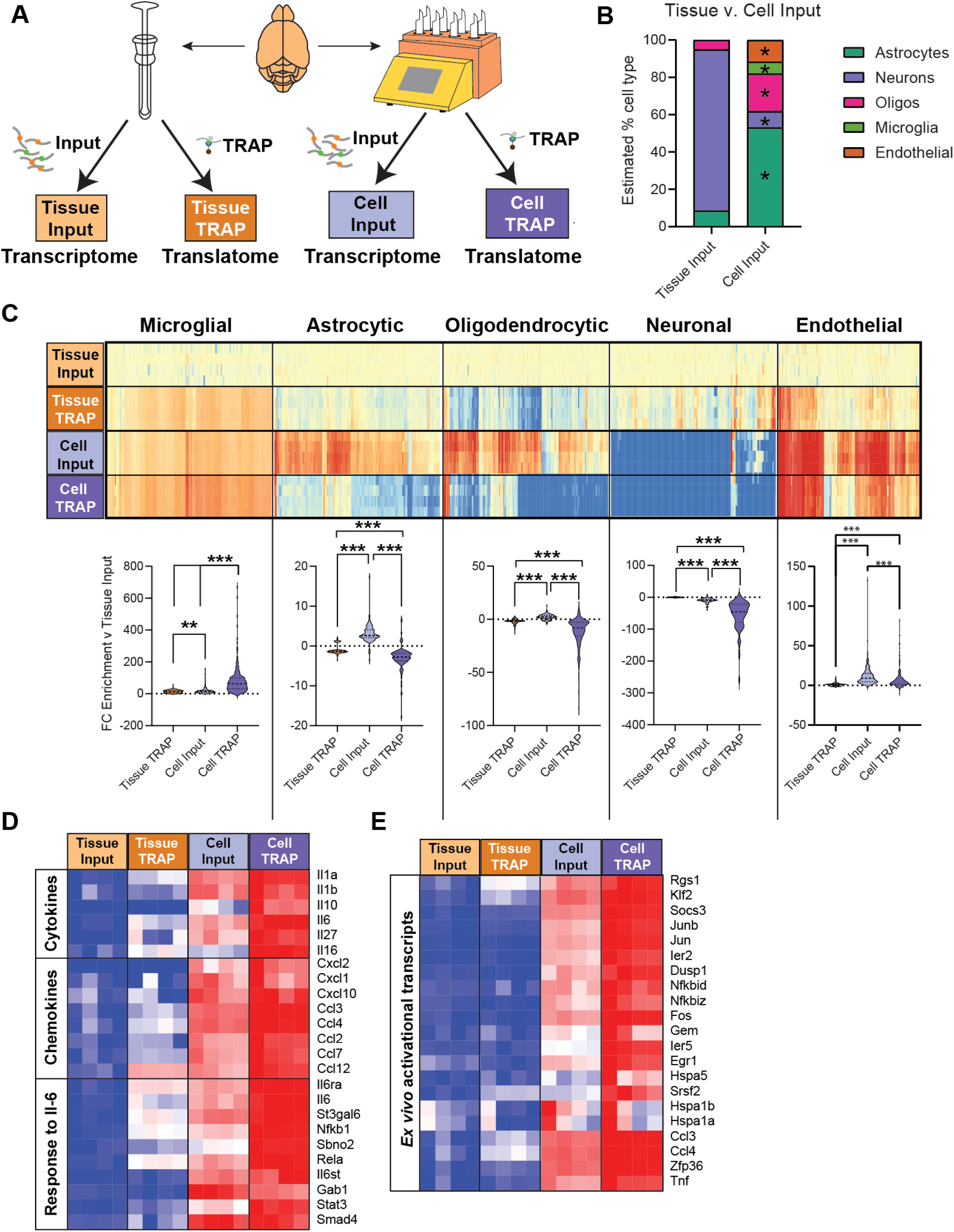
Changes in cellularity and *ex vivo* activation during cell preparation. A) Schematic of experimental design presented in this figure. B) CIBERSORTx cellularity estimates based on whole-transcriptome RNA-Seq from Tissue Input and Cell Input. C) Heatmap and violin plots of cell type-specific markers for microglia, astrocytes, oligodendrocytes, neurons, and endothelial cells (One-way ANOVA, Tukey’s post-hoc, *p<0.05, **p<0.01, ***p<0.001). D) Heatmap of inflammatory cytokines, chemokines, and response to Il-6 pathway genes. E) Heatmap of *ex vivo* activational genes identified in at least two previous studies (Ayata et al., 2018; Haimon et al., 2018; Marsh et al., 2020).

### Effect of transcriptional and translational inhibitors on *ex vivo* activational profiles following cell preparation

Recent studies have suggested that inclusion of transcriptional and/or translational inhibitors during cell preparation can prevent *ex vivo* activational confounds in microglia (Marsh et al., 2020) and other neuronal cell types (Wu et al., 2017). To test the effect of transcriptional and translational inhibitors on the transcriptome and translatome from cell suspension, we supplemented our cell preparation enzymatic mix with transcriptional and translations inhibitors: Actinomycin D, Triptolide, and Anisomycin. To assess the effect of transcriptional and translational inhibitors on the *ex vivo* activational state of microglia following cell preparation, the TRAP-isolated translatomes from tissue homogenate (Tissue-TRAP) was compared to TRAP from isolated cells without inhibitors (Cell TRAP – inhibitors) and with inhibitors (Cell TRAP + inhibitors) (**Figure 6A**). There were 121 genes that were higher in Cell TRAP -inhibitors compared to Tissue-TRAP. The majority (111/121=92%) of the genes that were induced during cell preparation were prevented by the addition of transcriptional and translational inhibitors during cell preparation (**Figure 6B**; **Supplemental Table 7**). Heatmap of the 111 *ex vivo* activational transcripts identified in **Figure 6B** shows addition of inhibitors during cell preparation partially ameliorates the *ex vivo* activational confounds induced by enzymatic and mechanical dissociation of brain tissue (**Figure 6C**; **Supplemental Table 7**). Top 10 biological processes over-represented in the 111 *ex vivo* activational genes prevented by the addition of inhibitors include: chemokine response, bacterial molecule response, and mechanical stimulus response, among others (**Figure 6D**; **Supplemental Table 7)**. Heatmap of response to mechanical stimulus pathway genes (**Figure 6E**; **Supplemental Table 7**) and response to chemokine (**Figure 6F**; **Supplemental Table 7**) show prevention of pathway induction with the use of transcription and translation inhibitors during cell preparation.

**Figure 6.**
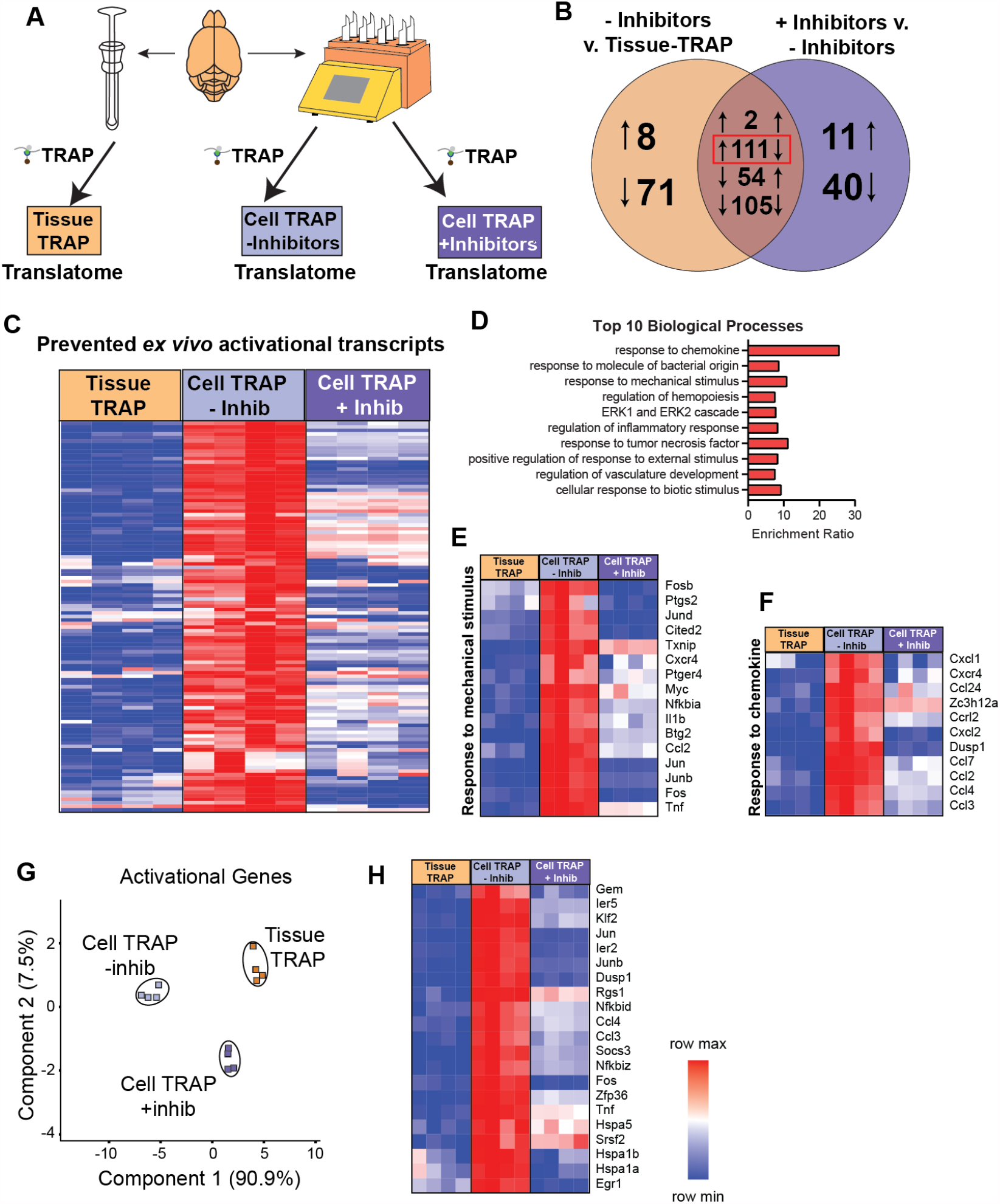
*Ex vivo* activational profiles with transcriptional and translational inhibitors. A) Schematic of experimental design presented in this figure. B) Differentially expressed genes were called between Cell-TRAP (+/-Inhibitors) and Tissue-TRAP. A subset of 111 genes were activated with cell preparation (Up in Cell-TRAP -Inhibitors v Tissue-TRAP) were decreased with the addition of inhibitors (Down in Cell-TRAP + Inhibitors v. Cell-TRAP – Inhibitors) (One-Way ANOVA, BHMTC, SNK FDR<0.1, |FC|>2). These 111 genes were classified as *ex vivo* activational transcripts prevented by the addition of inhibitors. C) Heatmap of the 111 *ex vivo* activational genes prevented by the addition of inhibitors. D) Top 10 biological processes over-representation in the 111 *ex vivo* activational genes prevented by the addition of inhibitors (hypergeometic test, BHMTC, FDR<0.05). E) Heatmap of response to mechanical stimulus biological process pathway. F) Heatmap of response to cytokine biological process pathway. G) PCA of 21 *ex vivo* activational genes identified in at least two previous studies (Ayata et al., 2018; Haimon et al., 2018; Marsh et al., 2020)

PCA of the 21 *ex vivo* activational transcripts identified in at least two of the three examined previous studies (Ayata et al., 2018; Haimon et al., 2018; Marsh et al., 2020) shows strong separation of Cell TRAP -inhibitors from Tissue-TRAP in the first component (90.9% explained variance). Addition of transcription and translation inhibitors during cell preparation migrated the Cell TRAP +inhibitors group closer to the Tissue-trap group in the first component (**Figure 6G**). Heatmap of the 21 *ex vivo* activational transcripts shows prevention of *ex vivo* activation with the addition of transcriptional and translational inhibitors (**Figure 6H**; **Supplemental Table 7**).

In summary, the addition of transcriptional and translational inhibitors during enzymatic and mechanical dissociation of brain tissue can prevent *ex vivo* activational artifacts in microglia.

### Effect of transcriptional and translational inhibitors on cell abundance following cell preparation

To verify that the prevention of *ex vivo* artifacts with the addition of inhibitors was not a result of altered cellularity during cell preparation, the Cell Input + Inhibitors was compared to Cell Input – Inhibitors and Tissue Input for changes in relative cell abundance by flow cytometry and transcriptomic analyses. Schematic of this workflow is represented in **Figure 7A**. Digital cytometry (CIBERSORTx) analysis on Tissue Input and Cell Input (+/-Inhibitors) showed differences in relative cell abundance between Cell Input (+/-Inhibitors) compared to Tissue Input (One-way ANOVA, Tukey’s post-hoc, *p<0.05). However, there were no significant differences in the relative cell-type proportions between Cell Input + Inhibitors compared to Cell Input – Inhibitors (**Figure 7B**; **Supplemental Table 8**). Next, we assessed the relative enrichment/depletion of microglial, astrocytic, oligodendrocytic, neuronal, and endothelial cell markers. Consistent with the results from **Figure 5**, we observed an overall enrichment of microglial, astrocytic, oligodendrocytic, and endothelial cell markers and depletion of neuronal cell markers in Cell Input (+/-Inhibitors) compared to Tissue Input. There was a small but significant difference in microglial and oligodendrocytic marker enrichment between Cell Input – Inhibitors and Cell Input + Inhibitors (Paired t-test, Bonferonni correction, *p<α=0.01) (**Figure 7C**; **Supplemental Table 8**). Flow cytometric analysis of cell type-specific markers revealed a small, but significant, difference in the relative abundance of oligodendrocytes with the addition of transcription and translation inhibitors as evidenced by the increase in O4+ singlets (Paired t-test, Bonferonni correction, *p<α=0.0125) (**Figure 7D**; **Supplemental Table 8**). In summary, the addition of transcriptional and translational inhibitors causes minimal changes in the relative abundance of cell types following cell preparation.

**Figure 7.**
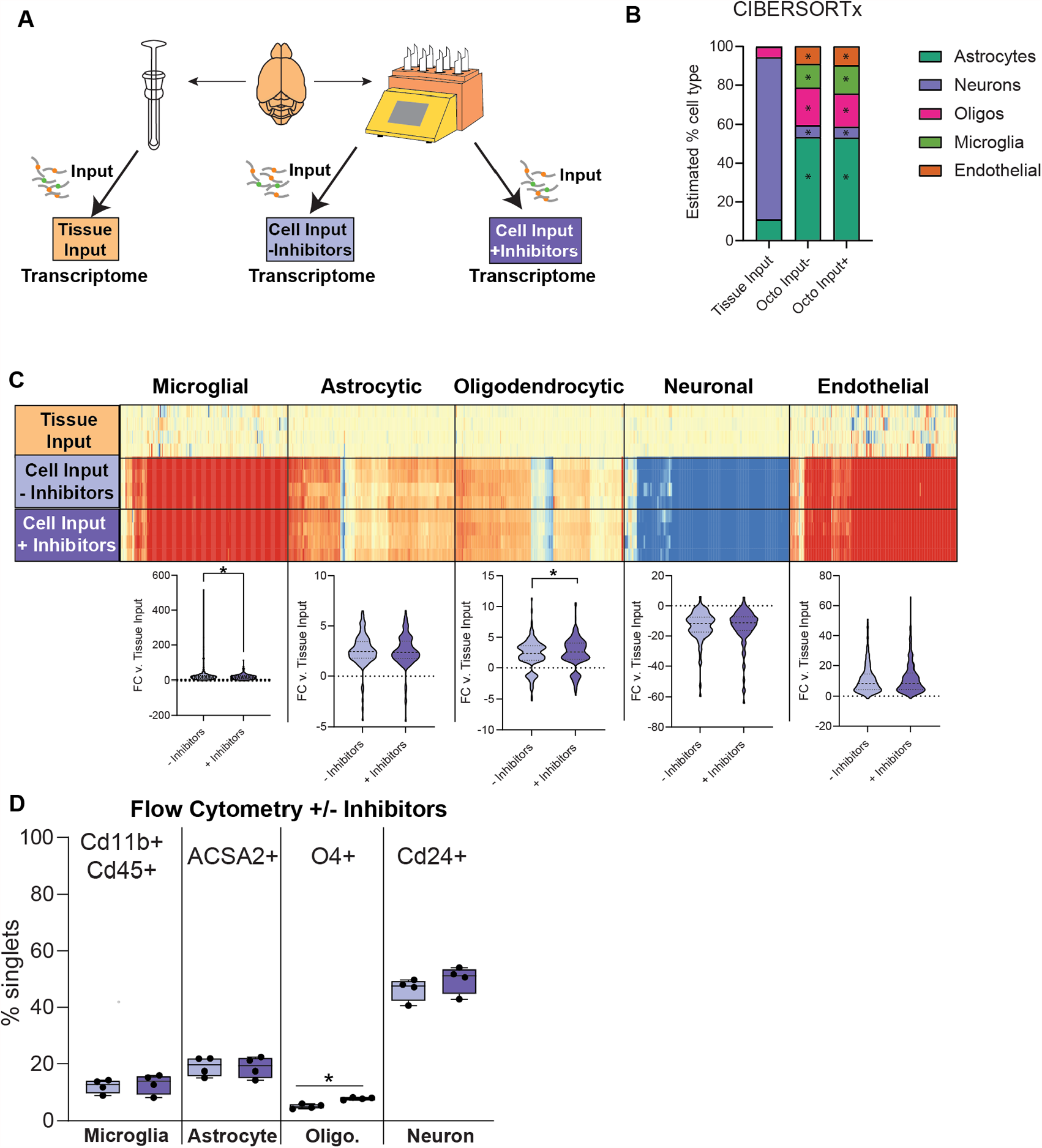
Effect of transcriptional and translational inhibitors on cell abundance. A) Schematic of experimental design presented in this figure. B) CIBERSORTx cellularity estimates based on whole-transcriptome RNA-Seq from Tissue Input and Cell Input (with and without inhibitors). There were significant differences in the proportions of each cell type between Cell Input +/-Inhibitors and Tissue Input (One-way ANOVA, Tukey’s post hoc, *p<0.05). However, there were no significant differences in relative cell abundances between Cell Inputs with the addition of transcriptional and translational inhibitors. C) Heatmap and violin plots of cell type-specific markers for microglia, astrocytes, oligodendrocytes, neurons, and endothelial cells (Paired t-test, Bonferonni correction, *p<α=0.01). D) Flow cytometry on cell type-specific markers for microglia (Cd11b+Cd45+), astrocytes (ACSA2+), and neurons (Cd24+) shows no significant differences in cell abundances with the addition of transcriptional and translational inhibitors. Flow cytometry on cell type-specific markers for oligodendrocytes (O4+) shows a small, but significant, increase in oligodendrocytes with the addition of transcriptional and translational inhibitors (Paired t-test, Bonferonni correction, *p<α=0.01).

## Discussion

Microglia have emerged as key players in brain disease, including age-related neuroinflammation and neurodegeneration (Colonna & Butovsky, 2017; Lana, Ugolini, Nosi, Wenk, & Giovannini, 2021). As a minority cell population in the brain, in depth microglial molecular and biochemical analyses benefit from enrichment strategies to provide cell-specific data (Okaty, Sugino, & Nelson, 2011). While *in vitro* cell culture models are useful for mechanistic studies, they fail to recapitulate the complexity of the nervous system milieu. As such, models and methods to ‘debulk’ microglia from brain tissue have become a focus in the field (Chucair-Elliott et al., 2020; McKinsey et al., 2020). The most common methods for isolating microglia include enzymatic and/or mechanical dissociation of brain tissue followed by immunolabeling with magnetic beads (Bordt et al., 2020) or fluorescent-conjugated antibodies for FACS-based enrichment (Bohlen, Bennett, & Bennett, 2019). As these methods have continued to evolve, quantitative comparisons of these methods are needed to aid decision making of what approaches to use in specific studies. As well, there are legitimate concerns that these isolation approaches introduce artifacts, especially in glial cell populations which by nature are sensitive to changes in their microenvironment (Wu et al., 2017). Determining the degree of *ex vivo* activational artifacts and how they may vary between isolation approaches, has been challenging because the field lacked a resting cell type-specific reference (absent of *ex vivo* activational confounds related to enzymatic/mechanical dissociation) as a comparator.

To address this barrier to progress, we used a ribosomal-tagging model (NuTRAP) in combination with a microglial-specific cre (Cx3cr1-cre/ERT2) to generate a microglial signature without the confounds of cell isolation. We then compared relative microglial enrichment and *ex vivo* activational artifacts between multiple MACS-and FACS-based cell isolation techniques. All sort methods were successful in isolating highly pure microglia, as evidenced by flow cytometric and transcriptomic analyses. Our MACS -FACS comparison is in line with previous findings on yield and speed (Pan & Wan, 2020). Magnetic bead-based isolation produced nearly as pure of a microglial population as FACS based approaches. The advantages of the magnetic bead isolation include the rapid isolation (<1hr), multiplexing 6 samples at a time, and least amount of instrumentation. The limitation of magnetic bead-based approaches is the single dimension of labeling as compared to FACS. Cartridge-based FACS is a new iteration of FACS and produced nearly equivalent cell purities as traditional cytometry-based FACS with greater cell yield. Another advantage of this cartridge-based approach is the self-contained nature of the system that does not produce aerosols thus not requiring biosafety containment (Roberts et al., 2021). Cytometry-based FACS produced the purest cell population and has the highest capabilities for dimensions of labeling. However, this approach is also the slowest and had the lowest cellular yield. A strict comparison of FACS methods relies on details of the gating strategies used and these can likely be tuned to emphasize highest purity or cell yield in both approaches. Combining MACS with cartridge-based FACS, to initially de-bulk microglia and then further purify, did not result in higher purity of microglia and the presence of the magnetic beads shifted signals in the FACS. Taken together, these data demonstrate that all of these approaches are valid for microglial isolation from brain and return highly pure cell populations that are suitable for molecular and biochemical analyses.

An unexpected finding of the analyses was the shift in cellularity caused by the cell suspension preparation. Neuronal cells and transcriptomic signals were depleted during cell preparation. While this has the effect of aiding microglial isolation by diminishing the majority neuronal cell population, this has profound effects on neuronal cell isolation studies. Outside of these studies, others have observed that alternative cell preparation methods cause less neuronal cell loss (Saxena et al., 2012). We did not test alternate cell preparation approaches such as these for their differential effects on microglial *ex vivo* activation and it remains possible that these different methods, through causing less cell death, could lead to less microglial activation. The large neuronal loss during cell preparation may contribute to microglia activation *ex vivo*.

Using TRAP isolation of the microglial translatome as a baseline measure of microglia, induction of *ex vivo* activational pathways occurred during enzymatic and mechanical cell preparation and was sustained during microglial isolation, independent of sort method. Compared to previous studies of these activational artifacts (Ayata et al., 2018; Haimon et al., 2018; Marsh et al., 2020) we found a common set of activational markers centered on immediate early genes. This unified set of markers can be used in future studies as markers of artifactual microglial cell activation.

The use of transcriptional and translational inhibitors (Marsh et al., 2020) during the cell preparation was investigated in the context of acutely isolating cells for immediate use in downstream molecular and biochemical analyses. Addition of transcription and translation inhibitors blocked much of the *ex vivo* activational artifacts without otherwise changing the cell phenotype. Whether this is a valid approach for cells that will subsequently be cultured was not assessed. Nonetheless, as studies also delve into microglial heterogeneity at the single cell level (Masuda, Sankowski, Staszewski, & Prinz, 2020; Stratoulias, Venero, Tremblay, & Joseph, 2019) inclusion of inhibitors in the preparation and enrichment of microglia can reduce artifacts in these studies as well (Marsh et al., 2020).

Collectively, our data demonstrate that a variety of microglial isolation methods can be used with equivalent results and tuned to the needs of the specific study. In addition, activational artifacts occur during cell isolation and can be prevented by inclusion of specific inhibitors early in the cell preparation protocol. The different cell sorting methods did not show additional activational effects or differences between methods, indicating that concerns over artifacts should not drive isolation method selection. Addition of transcriptional and translational inhibitors during cell preparation reduces *ex vivo* artifacts and is an easily implementable approach to avoid potential confounds.

## Acknowledgments

This work was supported by grants from the National Institutes of Health (NIH) P30AG050911, R01AG059430, R56AG067754, T32AG052363, F31AG064861, P30EY021725, Oklahoma Center for Adult Stem Cell Research (OCASCR), a program of the Oklahoma Tobacco Settlement Endowment Trust, BrightFocus Foundation (M2020207), and Presbyterian Health Foundation. This work was also supported in part by the MERIT award I01BX003906 and a Shared Equipment Evaluation Program (ShEEP) award ISIBX004797 from the United States (U.S.) Department of Veterans Affairs, Biomedical Laboratory Research and Development Service. The authors would also like to thank the Clinical Genomics Center (OMRF), Imaging Core Facility (OMRF) and DMEI (OUHSC), and Flow Cytometry and Cell Sorting Core Facility (OMRF) for assistance and instrument usage and Jan Brandel for assistance with illustrations. Some reagents and loan of a cytometer instrument were supplied at no cost by Miltyeni Biotec.

## Author contributions

Sarah R. Ocañas: first author, design of the work, execution of experiments, data acquisition, analysis, and interpretation, figure generation, manuscript writing and preparation

Kevin Pham: execution of experiments, data acquisition, analysis, and interpretation, figure generation, manuscript writing and preparation

Harris Blankenship: execution of experiments, data acquisition, analysis, and interpretation, manuscript writing and preparation

Adeline Machalinski: execution of experiments, data acquisition, analysis, and interpretation

Ana J. Chucair-Elliott: design of the work, execution of experiments, data acquisition, analysis, and interpretation, figure generation, manuscript writing and preparation

Willard M. Freeman: Corresponding author, conceptual design of the study, data analysis and interpretation, figure generation, manuscript writing, preparation, and submission.

---

**Figure S1.**
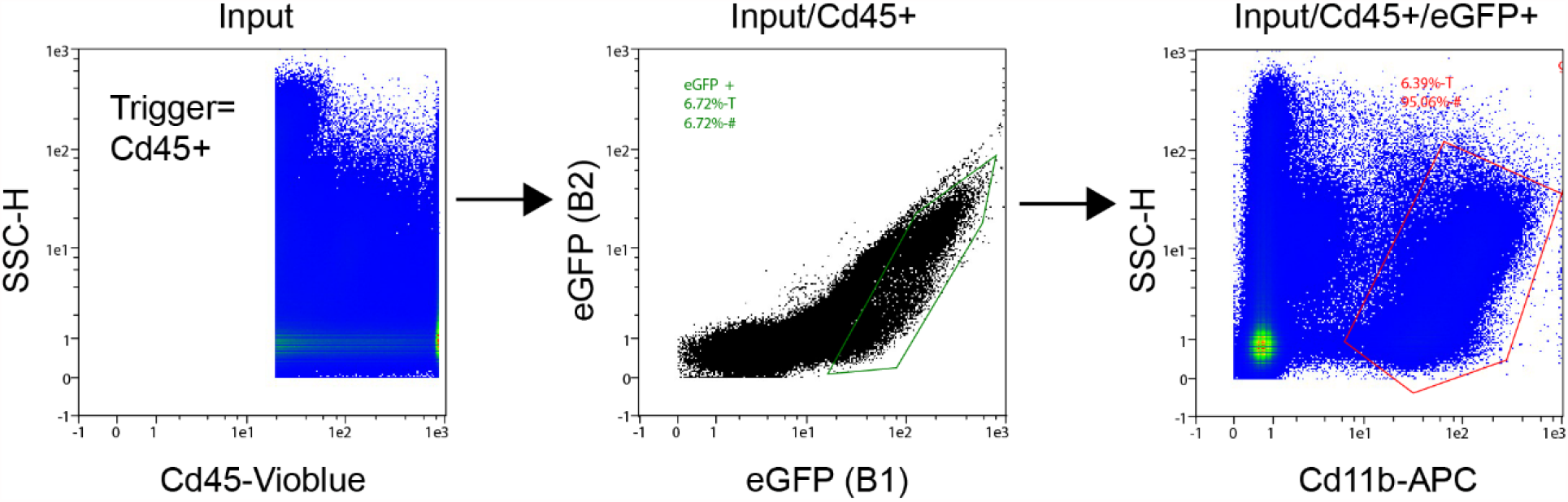
Cartidge-based FACS gating strategy for microglial sorting on Miltenyi Biotec MACSQuant Tyto.

**Figure S2.**
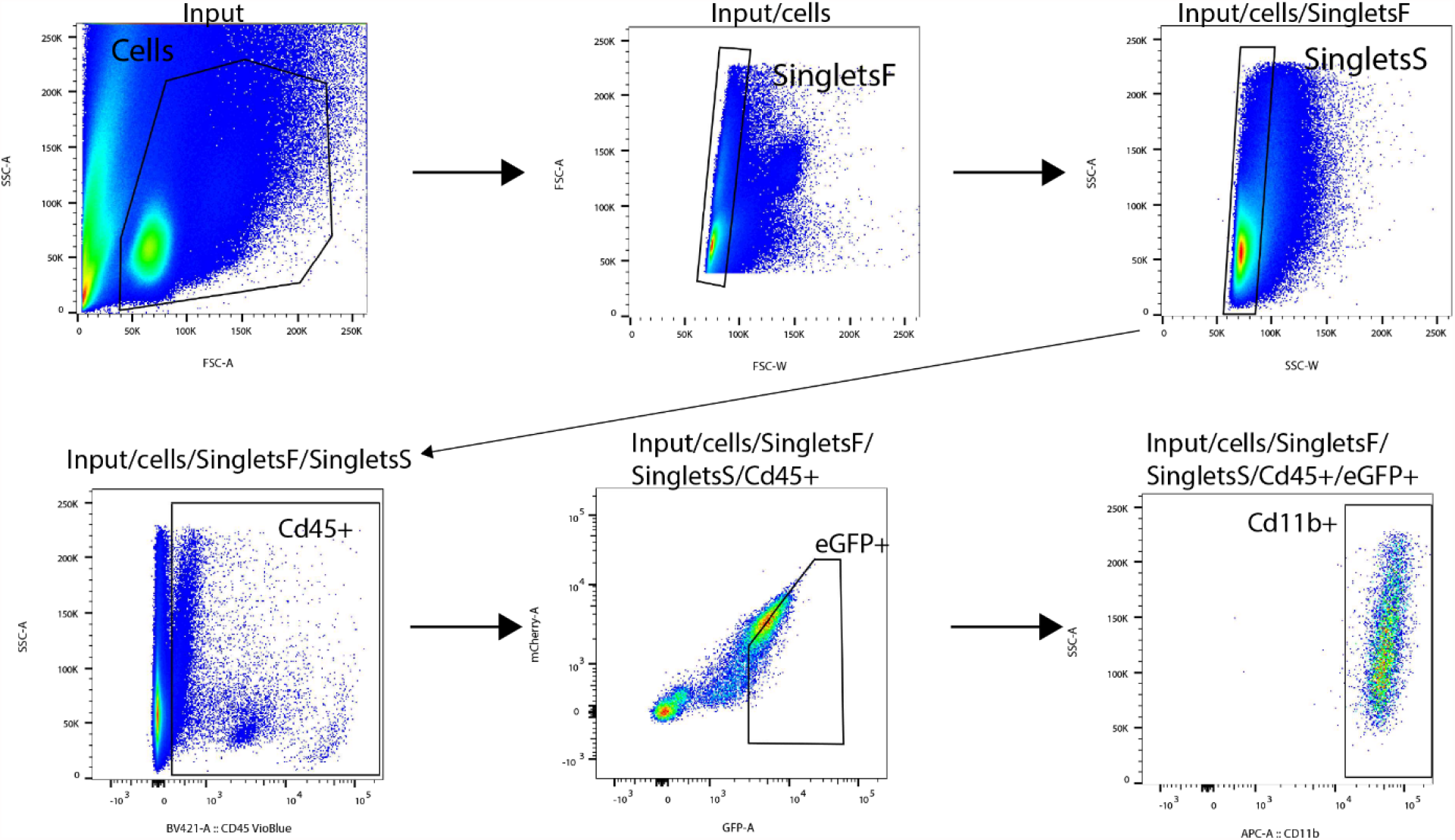
Cytometer-based FACS gating strategy for microglial sorting on FACSAria.

**Figure S3.**
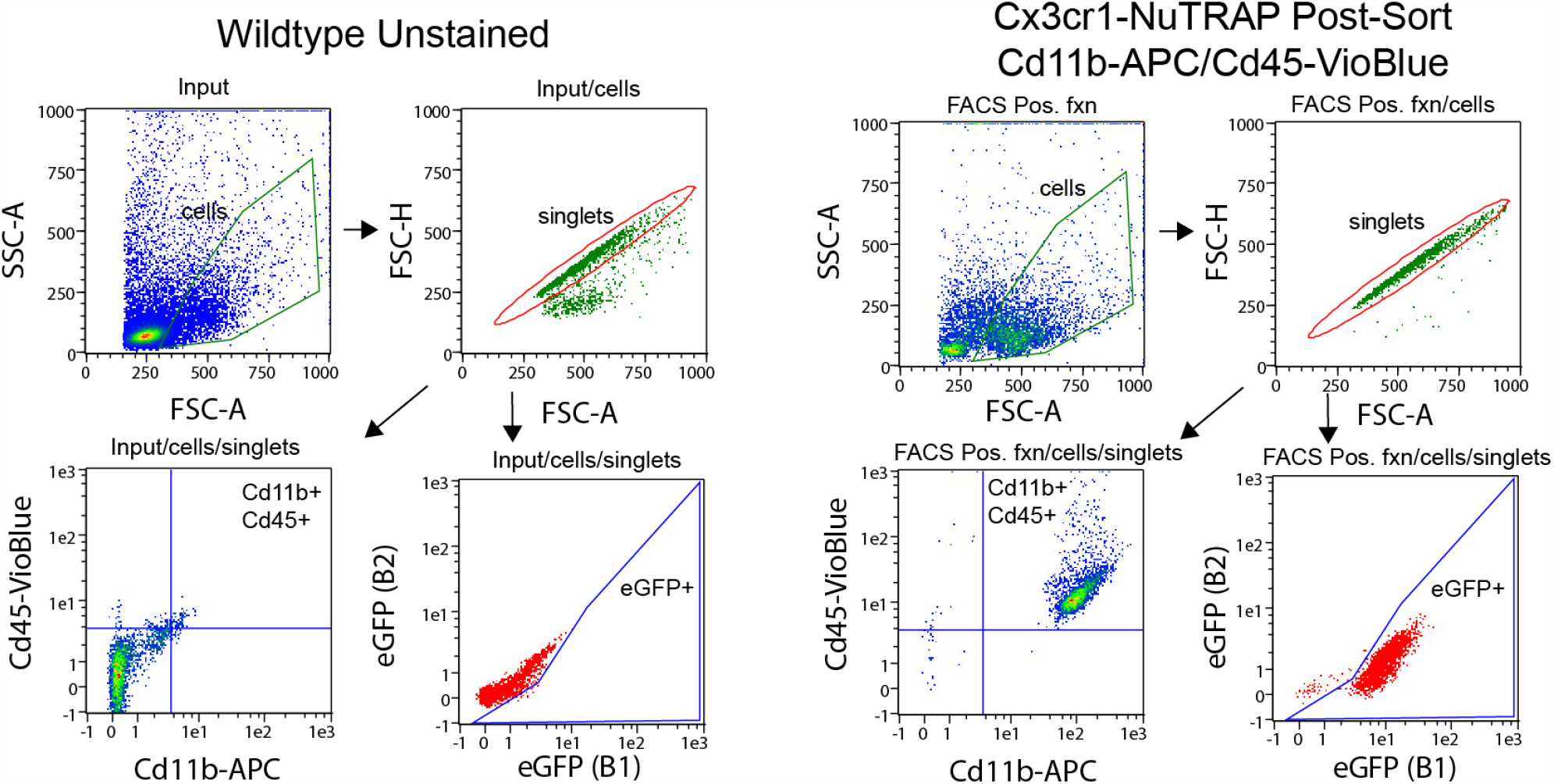
Gating strategy for assessment of microglial purity by different sort methods.

**Figure S4.**
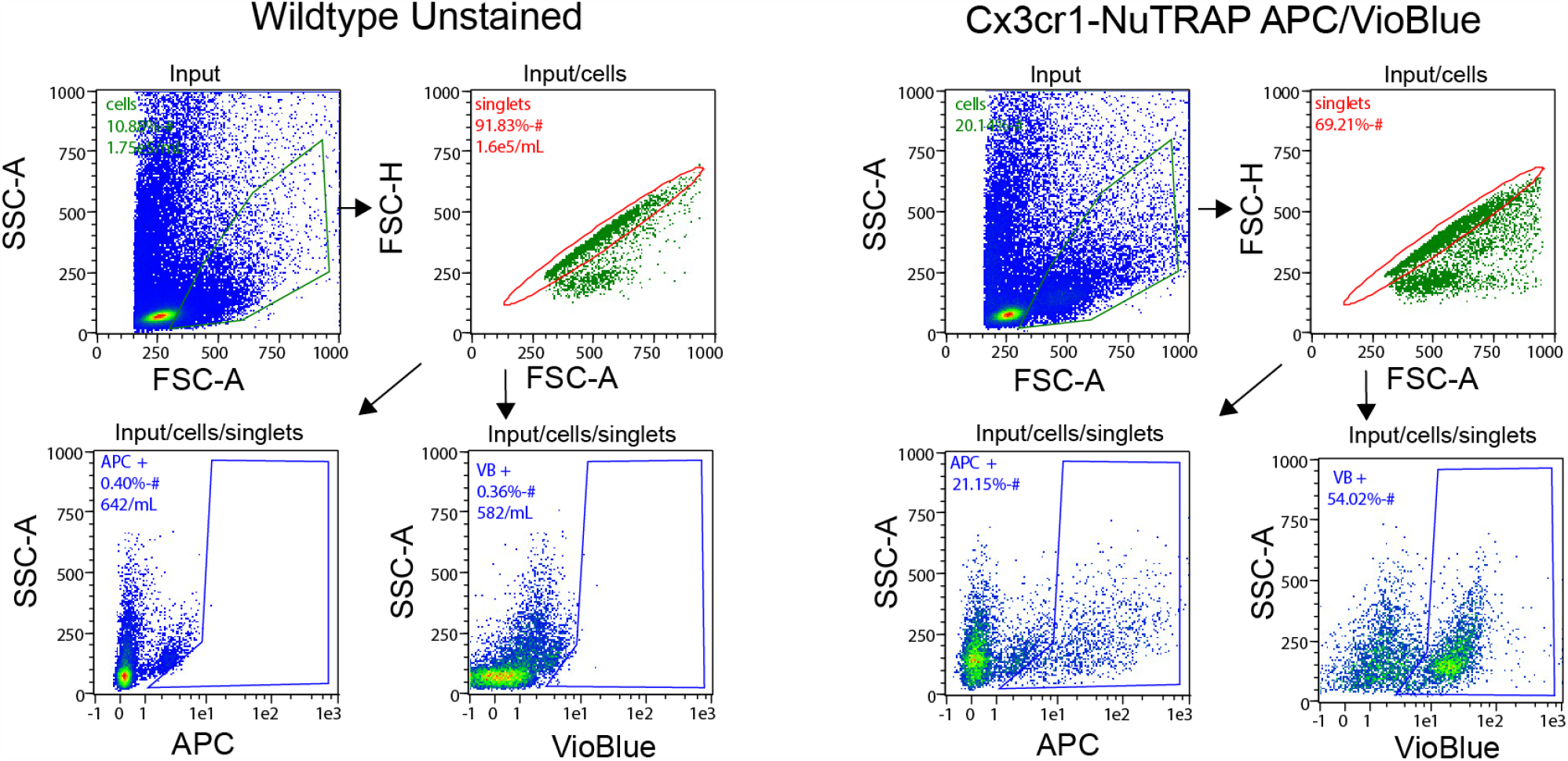
Gating strategy for assessment of cellularity.

**Figure S5.**
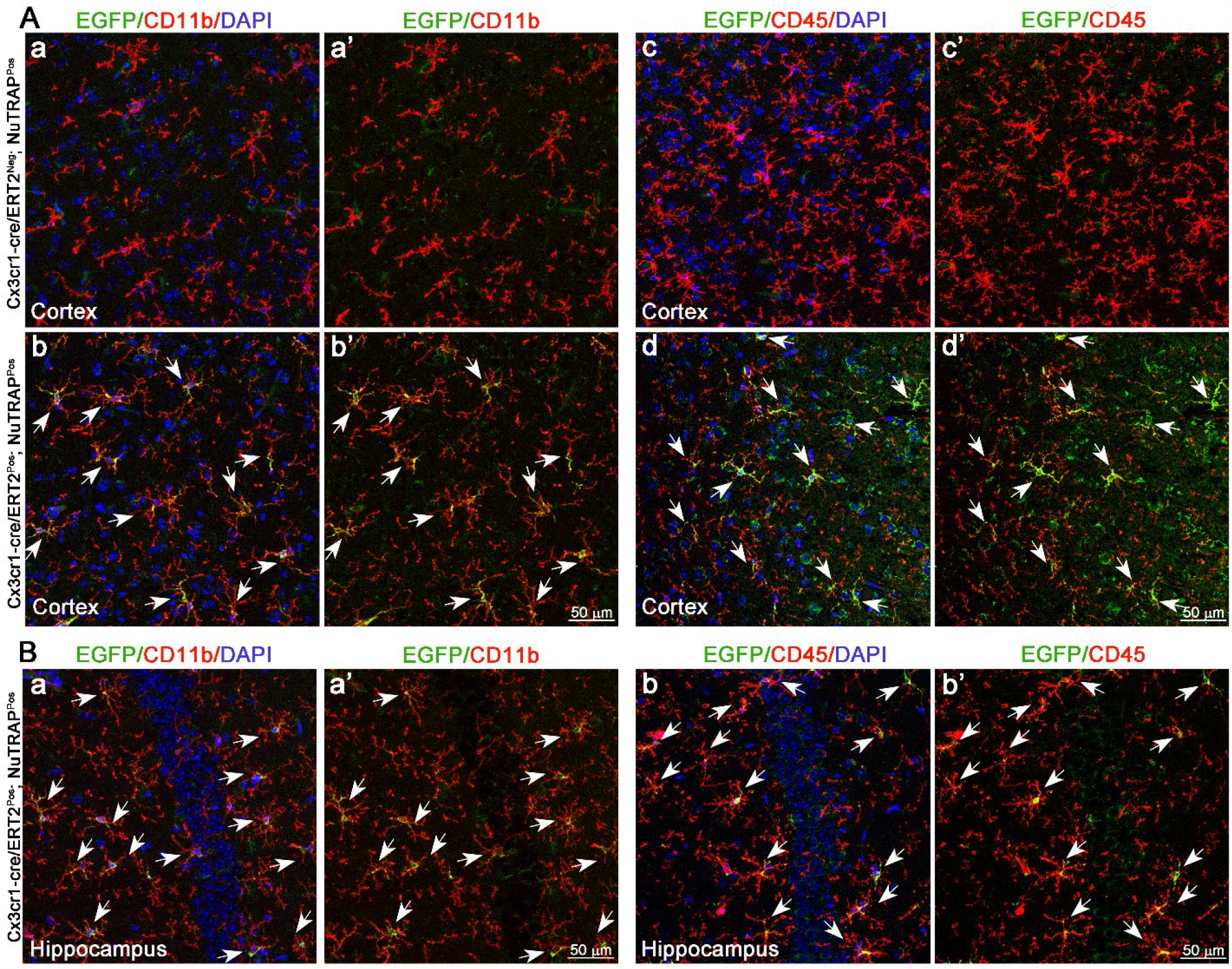
Validation of microglial identity of recombined cells in the Cx3cr1-NuTRAP brain. Two months after Tam treatment, brains were harvested from Cx3cr1-NuTRAP and cre negative NuTRAP^+^ (control) mice for immunohistochemistry (IHC). **A**. Representative confocal fluorescent microscopy images of sagittal brain sections captured in the cortex show EGFP expression (green signal) in cells that co-expressed CD11b (red signal, **a-a’-b-b’**) and CD45 (red signal, **c-c’-d-d’**) in Cx3cr1-NuTRAP brains but not in the cre negative counterparts (n=2/ group).**B**. Representative confocal fluorescent microscopy images captured in the hippocampus show EGFP expression (green signal) in cells that co-expressed CD11b **(a-a’)** and CD45 **(b-b’)** in Cx3cr1-NuTRAP brains DAPI: nuclear counterstain. Scale bar: 50 µm

