## Supplemental Figures for "Minimizing the *ex vivo* confounds of cell-isolation techniques on transcriptomic -profiles of purified microglia"

**Affiliations:** [1]Genes & Human Disease Program, Oklahoma Medical Research Foundation, Oklahoma City, OK USA, [2]Department of Physiology, University of Oklahoma Health Sciences Center, Oklahoma City, OK USA, [3]Oklahoma City Veterans Affairs Medical Center, Oklahoma City, OK USA, [4]Department of Biochemistry and Molecular Biology, University of Oklahoma Health Sciences Center, Oklahoma City, OK USA

**To whom correspondence should be addressed:** [*]Willard M. Freeman, Genes & Human Disease Program, Oklahoma Medical Research Foundation, 825 NE 13th Street, Oklahoma City, OK 73104, USA.

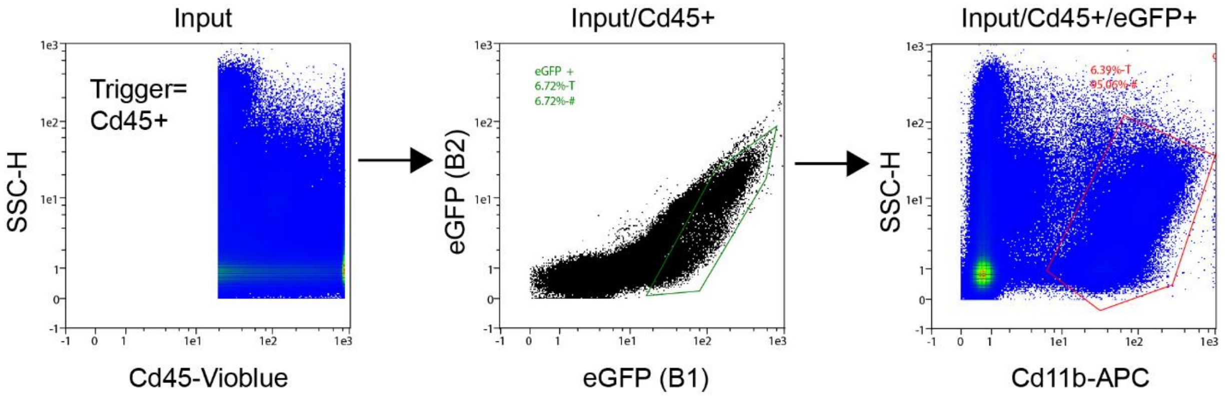


**Figure S1.** Cartidge-based FACS gating strategy for microglial sorting on Miltenyi Biotec MACSQuant Tyto.

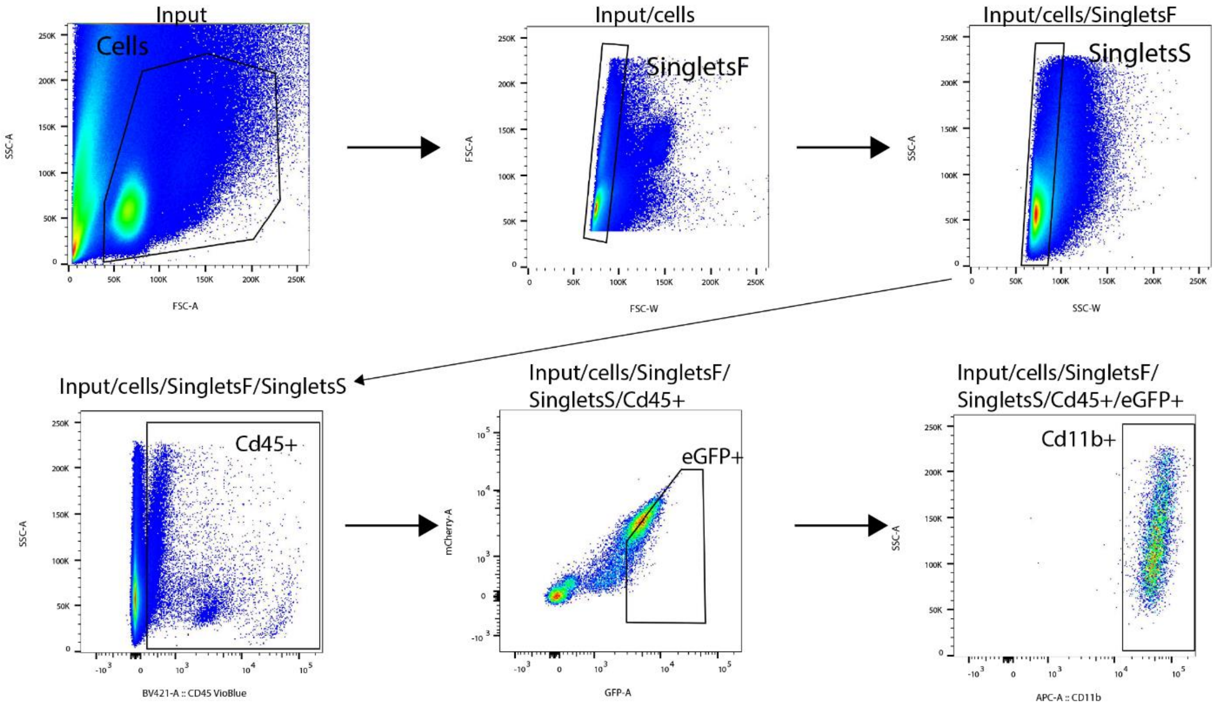


**Figure S2.** Cytometer-based FACS gating strategy for microglial sorting on FACSAria.

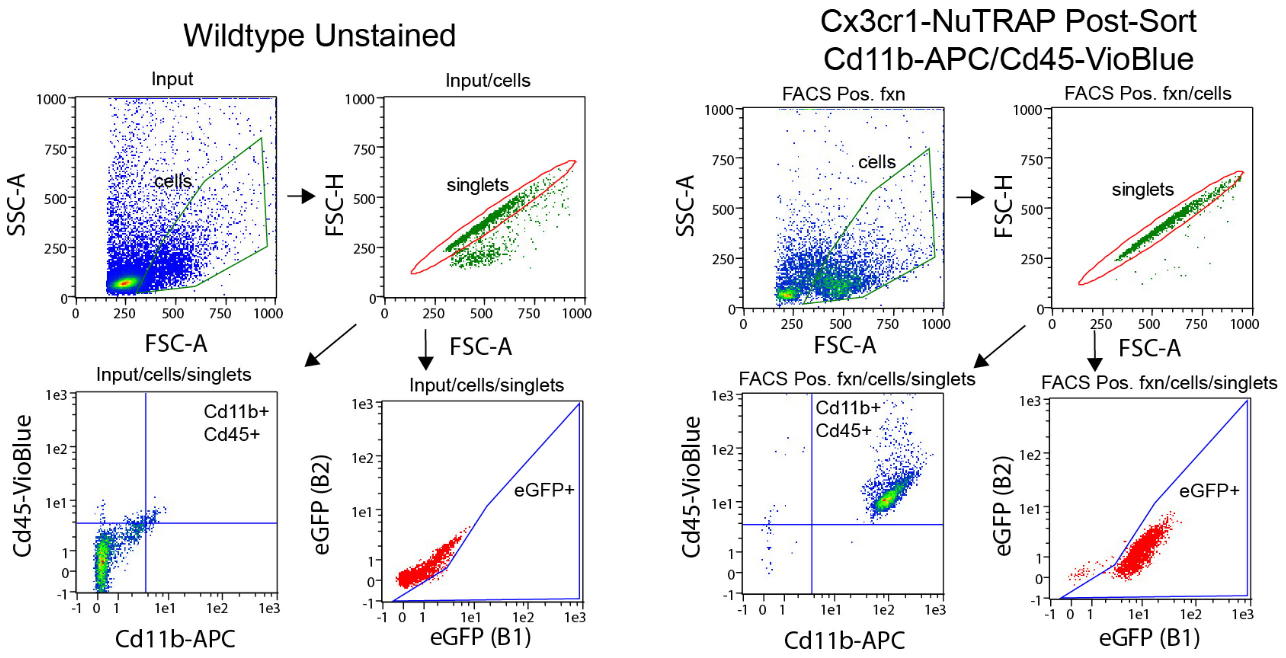


**Figure S3**. Gating strategy for assessment of microglial purity by different sort methods.

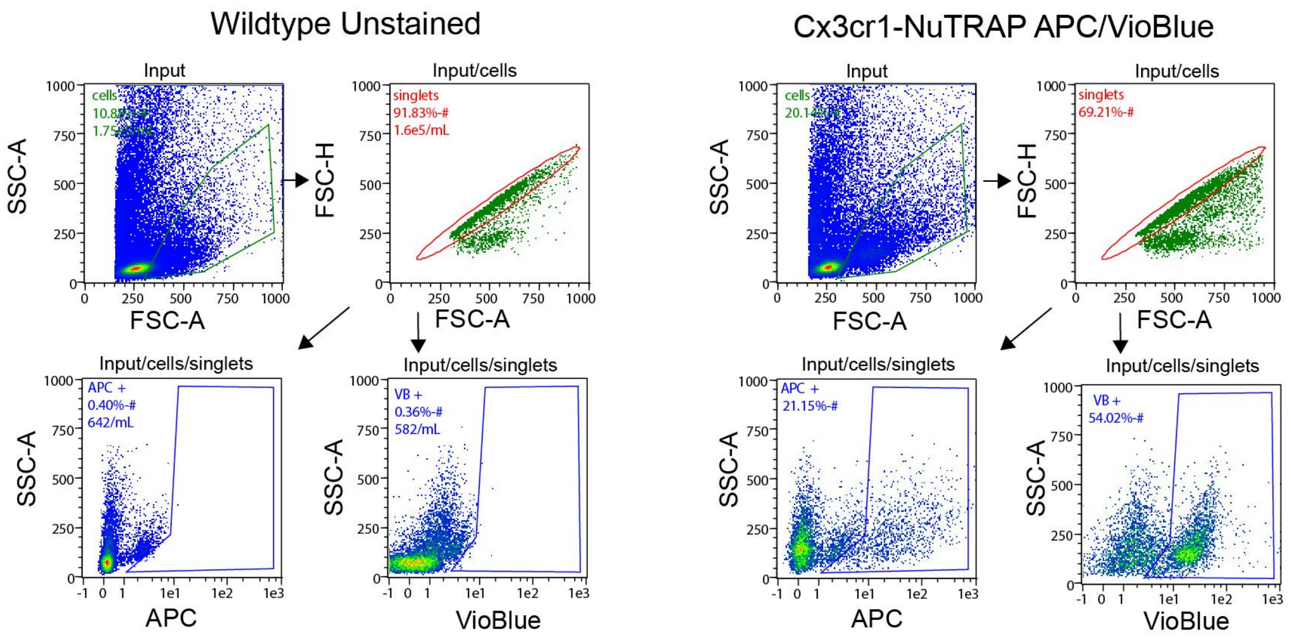


**Figure S4**. Gating strategy for assessment of cellularity.

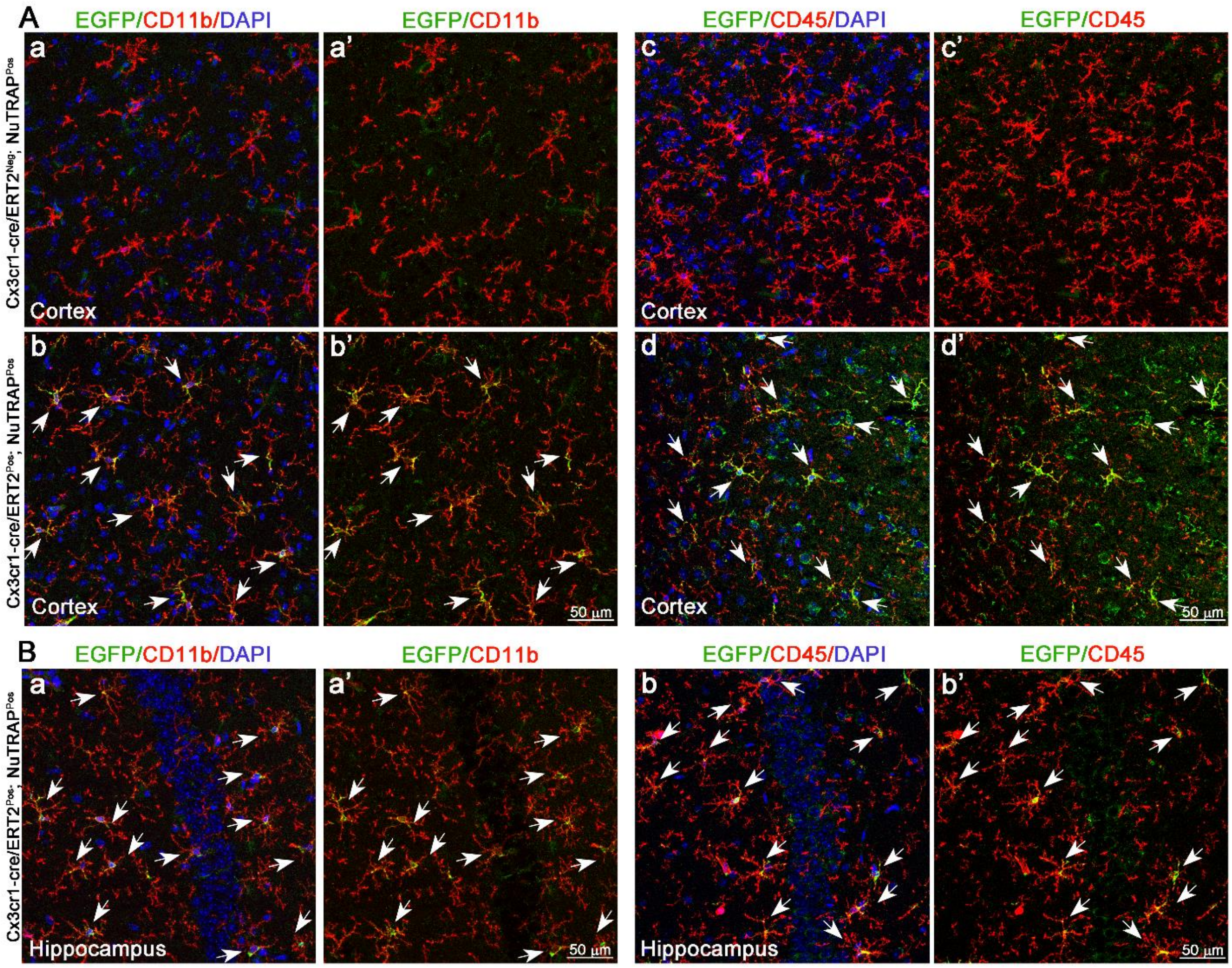


**Figure S5. Validation of microglial identity of recombined cells in the Cx3cr1-NuTRAP brain.** Two months after Tam treatment, brains were harvested from Cx3cr1-NuTRAP and cre negative NuTRAP$^{+}$ (control) mice for immunohistochemistry (IHC). **A.** Representative confocal fluorescent microscopy images of sagittal brain sections captured in the cortex show EGFP expression (green signal) in cells that co-expressed CD11b (red signal, **a-a'-b-b'**) and CD45 (red signal, **c-c'-d-d'**) in Cx3cr1-NuTRAP brains but not in the cre negative counterparts (n=2/ group). **B.** Representative confocal fluorescent microscopy images captured in the hippocampus show EGFP expression (green signal) in cells that co-expressed CD11b **(a-a')** and CD45 **(b-b')** in Cx3cr1-NuTRAP brains DAPI: nuclear counterstain. Scale bar: 50 µm
